# Pesticide chemical leads inhibiting protein-protein interactions

**DOI:** 10.1101/2024.02.03.578754

**Authors:** Rotem Shelly Ben Shushan, Elad Cohen, Noam Ben Naim, Eytan Amram, Jonathan Gressel, Dotan Peleg, Nesly Dotan, Itai Bloch, Maayan Gal

## Abstract

Pesticides, especially herbicides, have revolutionized agriculture by providing energy-efficient solutions for pest control that replaces labor-intensive cultivation methods. However, the widespread evolution of pesticide resistance poses a significant challenge to current agriculture. Most pesticides function by binding to specific pockets on target enzymes, enabling a single mutation to confer resistance. An alternative approach is the disruption of protein-protein interactions (PPI), thus for resistance to occur, it requires complementary mutations on both interacting partners. Despite extensive efforts, no herbicides with new modes of action have been commercialized for decades. Thus, we focused on the discovery and design of small molecule inhibitors that target the interface of the PPI complex of O-acetylserine sulfhydrylase (OASS) and serine acetyltransferase (SAT), key plant enzymes involved in the biosynthesis of the essential amino acid cysteine. Using in silico filtering techniques on a virtual library of 30 million small molecules, we identified initial hits capable of binding OASS and interfering with its interaction with a peptide derived from SAT. Subsequently, we conducted chemical optimizations to evaluate biophysical enzyme disruption, followed by cellular and in-planta activity in plants. These new compounds described herein can serve as promising starting points for further optimization as herbicides acting on a new mode of action.

## Introduction

### Resistance and no new modes of action decrease food security

The high-yield crop production necessary for global food security relies on effective weed management, which is dependent on the application of herbicides^1^. However, due to the heavy use of existing herbicides, resistance has evolved at an accelerated pace in the last few years^2^ and large cropping areas are now covered with one or more weeds of >500 species that have evolved resistance to one or now usually more herbicides with different modes of action^3^. The problem is exacerbated by the deregistration of many herbicides, such that fewer herbicides are available to deal with resistance and no new modes of action have been commercialized for decades, although a few appear to be in the pipeline. Farmers have had to revert to soil structure-disrupting, erosion-promoting, and mechanical cultivation, which is apparent in the increase in fossil fuel usage, which had been declining due to the increased use of herbicides^4^. Thus, there is an urgent need to develop new resistance-recalcitrant herbicides, acting by new modes of action that inhibit novel protein targets.

### Inhibiting protein-protein interactions: bringing vast new target possibilities

Most marketed pesticides act by binding to discrete enzymatic pockets, allowing a single mutation in the binding pocket to exclude the pesticide and resistance to evolve quickly. Various biochemical reactions within the cell require two or more peptides or enzymes to be in close proximity, i.e., the enzymatic activity requires protein-protein interactions (PPI). PPIs have an essential role in all layers of cellular activities and thus, disrupting PPIs opens the path to the discovery of pesticides to which resistance has not yet evolved. Moreover, as the number of PPIs (i.e., the interactome) is much larger than the number of single genes, there is a huge number of yet unexplored protein interfaces that can be targeted^5^. Targeting PPI interfaces can deal with two critical problems; mammalian toxicity and the evolution of resistance. Targeting PPIs that have no vertebrate homologies suggests an excellent safety profile. In addition, a single mutation that precludes pesticide binding to a PPI ‘hot spot’ would be lethal to the pest if the two proteins could not bond without a simultaneous complementary mutation on the interacting protein. The frequency of two simultaneous mutations is the compounded frequency of each event, an extremely low number.

PPIs had been considered to be “undruggable” as such interfaces are typically relatively large (more than 800 Å in diameter) and shallow, a challenge for the discovery of small binding molecules^6,7^. PPI disruptor discovery had been hampered by the requirement for *a-priori* structural data, which were not abundant, especially when at least one of the interaction partners is an intrinsically disordered protein^8,9^. Computer learning and modeling changed the situation allowing one to discern binding ‘hot-spots’ on the shallow PPI interface and further allowing the determining parameters of structures of small molecules that disrupt the PPI. This approach has been used by the pharmaceutical industry to design new drugs^10^.

Interfering with the PPIs involved in the biosynthesis of essential amino acids is an attractive approach for the design of new herbicides^11^. Cysteine has a significant role in plant growth and development as it is the primary source of the sulfur moiety in sulfur-containing compounds. The final reaction in the cysteine biosynthesis pathway involves two sequential reactions on interacting proteins. The first is catalyzed by serine acetyltransferase (SAT, EC 2.3.1.30), which synthesizes the intermediary product, O-acetyl serine (OAS), from acetyl-CoA and serine. The coupled reaction is catalyzed by O-acetylserine sulfhydrylase (OASS, EC 2.5.1.47), which incorporates sulfide to OAS producing L-cysteine. The C-terminus tail of SAT binding to OASS is essential for cysteine biosynthesis in plants and bacteria^12,13^. The SAT/OASS complex has a 3:2 stoichiometry, where the C-terminal region of three SAT subunits bind to the OASS homodimer^14-16^. SAT activity within the complex is negatively regulated by elevated cysteine and OAS concentrations, suggesting for the sensitivity of the complex to altering cellular conditions^17-19^. The SAT C-terminal peptide is seen anchored within the OASS active site in the crystal structure of Arabidopsis, where its highly conserved residue I314 carboxylate interacts with OASS amino acids Q147, T78, N77 and T74, and its hydrophobic side chain located towards the pyridoxal phosphate cofactor in a hydrophobic pocket (Fig. 1a-c). These highly conserved interactions are critical for complex formation, and mutations within this binding region abolished the OASS-SAT interaction^20^. The essentiality of SAT/OASS pathway was further validated by showing that OASS inhibitors reversed bacterial resistance to antibiotics, and that SAT inhibition by a small molecule hampered bacterial growth^20-22^. Based on the rational that PPI disruption can be a basis for pesticide discovery, and the essential role of the OASS-SAT interaction in cysteine biosynthesis, we sought to discover and develop new small molecules capable of interfering with this interaction *in-planta* as herbicide leads.

**Fig. 1.**
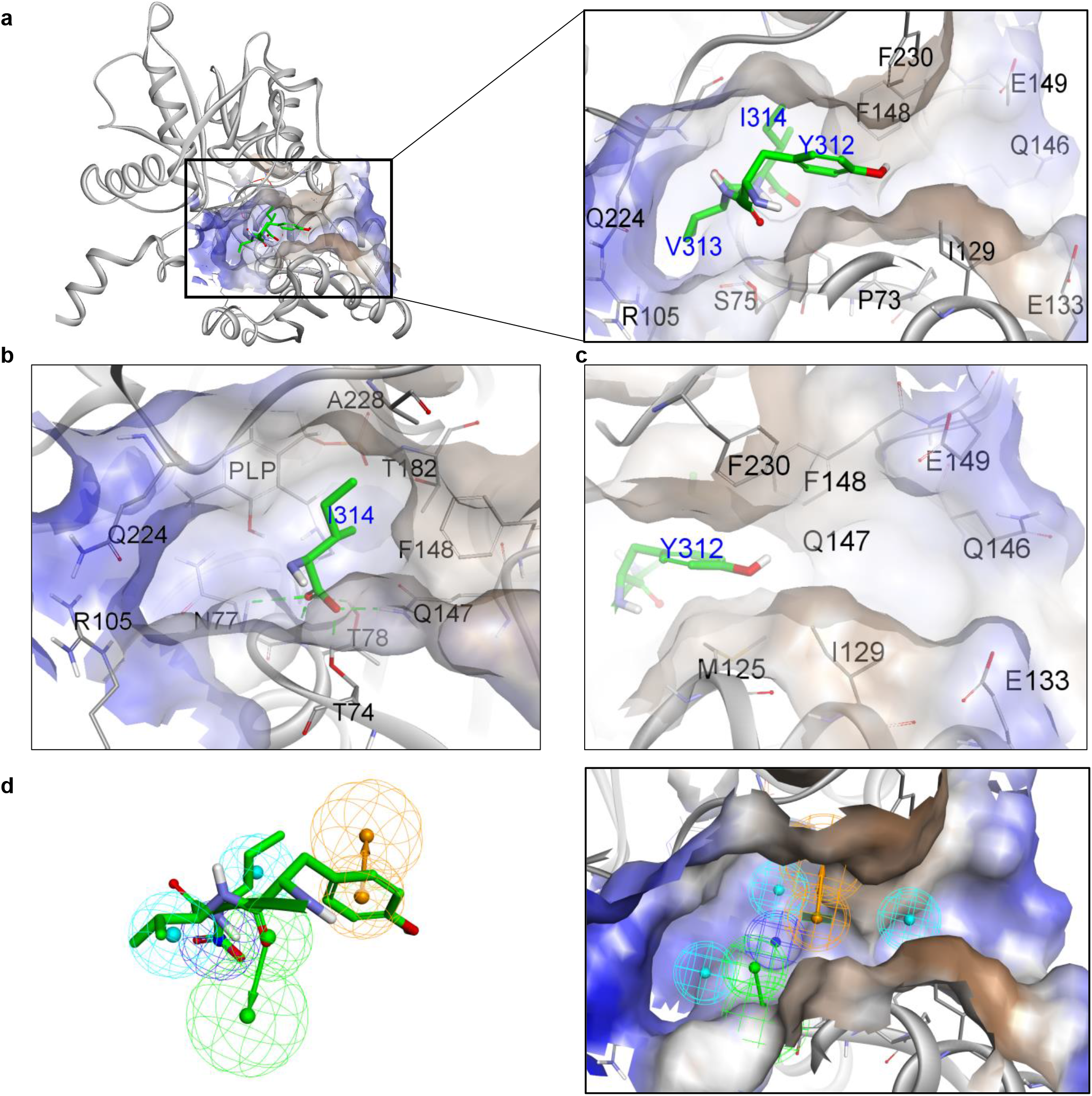
OASS Binding site and pharmacophore constraints. **a**, Structural model of OASS bound to SAT peptide YVI C-terminus residues based on protein data bank 2ISQ. **b**, Zoom in on SAT I314 and its interaction with OASS residues. **c**, Zoom in at a different view angle on SAT Y312 and its interaction with OASS residues. The OASS residues contributing to the interaction with SAT and/or defining the putative small molecule binding pocket are indicated. **d**, Illustration of the pharmacophore constraints that were defined on the YVI residues based on the model in **a**. The hydrophobic feature is indicated as a light blue sphere, hydrogen bond acceptor as a green sphere, negative charge as a blue sphere, and aromatic ring as an orange sphere.

## Results

### Filtering virtual libraries

We constructed a virtual library of about 30 million small organic compounds derived from various commercial sources to screen for small molecules with the potential of binding to OASS. We embedded a list of physio-chemical properties of known herbicides according to the ‘Tice rules’, a set of simple molecular criteria that have been derived from the analysis of hundreds of herbicides^23^. The criteria that were used to filter the virtual library are shown in Table 1. This step reduced the library size to 900,000 molecules. Although the molecular weight of some of the most used herbicides is <200 Da, to balance between the required physio-chemical features to inhibit PPI interfaces and considering that yet many herbicides are > 300 Da ^24^, we defined molecular weight of 300-600 Da.

**Table 1.** Author-defined physio-chemical properties used to filter the initial molecular library.

| Physio-chemical property | Min | Max |
| --- | --- | --- |
| Number of non-hydrogen atoms | 15 | 40 |
| Number of hydrophobic functional groups | 1 | 2 |
| Number of aromatic rings | 1 | 3 |
| Number of non-aromatic rings | 0 | 4 |
| Number of H acceptors | 2 | 7 |
| Number of H donors | 0 | 3 |
| Number of negative functional groups | 1 | 2 |
| Number of positive functional groups | 0 | 1 |
| Number of rotatable bonds | 2 | 8 |
| Log*P* (calculated or empiric) | 0.5 | 5.5 |
| Molecular Weight (Da) | 300 | 600 |

### Binding epitope and pharmacophore constraints to further filter potential leads

We then defined a set of chemical requirements and preferred characteristics for the binding of small molecules to the OASS interface. A thorough structural analysis of OASS was performed based on published X-ray structures of the enzyme with different ligands^20,25^. This step defined a potential small molecule binding site located in the vicinity of the OASS pyridoxal phosphate co-factor. The SAT binding site on the surface of OASS with the three C-terminus residues YVI is depicted in Fig. 1a. The site includes a hydrophobic pocket between F148, A228 and T182, followed by a hydrophilic pocket between Q147, T78, N77, Q224 and R105 (Fig. 1b). This latter is linked by a hydrophobic channel between F230, F148, M125 and I129 to a secondary hydrophilic pocket between E149, Q146 and E133 which is not occupied by the SAT C-terminal (Fig. 1c). We then constructed the binding pharmacophore, a set of structural constraints facilitating additional filtering of the small molecules’ library. These constraints were defined based on the analysis of the crystal structure of the Arabidopsis OASS/SAT interaction^20^, bacterial OASS structures with small molecule and peptide binders to OASS^25-28^. The pharmacophore volume constraints that were defined based on all the analyzed structures and marked on the YVI residues of SAT are illustrated in Fig. 1d.

The process described above resulted in a list of structural determinants, which constitute the full description of the pharmacophore (Table 2). The spatial XYZ coordinates in Table 2 dictate the location of a particular functional group or moiety of the potential molecule, followed by the radius that defines the sphere within which the functional group is to be placed. The center of the sphere is presented in angstrom (Å) on a cartesian coordinate system.

**Table 2.** Pharmacophore constraints.

| Name | Notation | X<br>(Å) | Y<br>(Å) | Z<br>(Å) | Radius<br>(Å) |
| --- | --- | --- | --- | --- | --- |
| Hydrophobic_1 | HD1 | 78.51 | 47.01 | -12.92 | 1.6 |
| Aromatic ring | AR1 | 77.12 | 51.92 | -10.61 | 1.6 |
| Aromatic ring projection | A1<sub>P</sub> | 80.05 | 51.32 | -10.93 | 2.2 |
| Negative atom | NC1 | 75.93 | 45.12 | -12.53 | 1.7 |
| H-bond acceptor | HA1 | 74.18 | 50.71 | -13.04 | 1.6 |
| H-bond acceptor projection | HA1<sub>P</sub> | 71.8 | 49.22 | -11.99 | 2.2 |
| Hydrophobic_2 | HD2 | 78.26 | 49.86 | -6.08 | 1.6 |

### Docking of selected molecules with OASS

The ∼400,000 molecules that passed through the initial pharmacophore constraints, were further evaluated by multiple rounds of molecular docking. At the end of the docking process, each molecule was assigned three orthogonal scores: 1. Pharmacophore fit score - determines the alignment of each molecule with the defined pharmacophore; 2. Docking score - evaluates the molecule torsion strain and type and the total number of interactions with OASS; 3. Pose score - evaluates the molecule spatial fitting relative to the interactions we defined as preferable between the molecule and OASS. Using this step, 160,000 molecules were selected and were further evaluated by additional orthogonal scoring functions. These were normalized and summed to a consensus score from which a final list of the top 20,000 molecules was constructed and arranged into 2000 clusters, which were analyzed manually. The final selection ensured that all molecules have a hydrophobic group where the I314 side chain is located, have a negatively charged group that makes hydrogen bonds with Q147, T78 and/or N77, as well as having an aromatic ring located near the place of SAT Y312 which serves as a linker to the secondary OASS pocket. We also validated the existence of optional hydrogen bonds with S75, Q224, Q146, E72, E133 or E149. This resulted in a final list of ∼180 molecules for in vitro evaluation for their ability to compete with OASS/SAT binding.

### Screening the in-vitro binding of selected virtual hits by fluorescence polarization

Each molecule was dissolved in DMSO (dimethyl sulfoxide) to a final concentration of 50 mM and individually tested for their ability to interfere with the OASS/SAT binding by single-dose fluorescence polarization (FP). The FP binding assay used recombinantly expressed OASS interacting with fluorescently labeled SAT-derived peptide. The excited fluorescent probe (i.e., SAT peptide) bound to a large molecular weight protein (i.e., OASS) emits light with a degree of polarization that is inversely proportional to the rate of molecular rotation. This fluorescence property is exploited to measure the interaction of a small labeled ligand with a larger protein and provides a basis for direct and competition binding assays^29,30^. Herein, we relied on the known SAT-derived peptide YLTEWSDYVI (SATp) conjugated with fluorescein isothiocyanate (FITC). Screening of the molecules binding to OASS and competing with FITC-SATp was executed by incubation of OASS with 50 μM of each molecule and 200 nM FITC-SATp and readout of polarization values. The latter were normalized based on milli polarization values (mP) of the free peptide (0% complex) and the FITC-SATp/OASS complex with no inhibitor (100% complex). The normalized mP values of the different compounds arranged in descending order is shown in Fig. 2a. This initial hit (red color) showed more than 50% reduction of the mP value, suggesting its effective inhibition of the complex formation. To validate the screening results, this molecule (PJ1) was tested in a full dose-response experiments, yielding an IC_50_ of 34 μM (Fig. 2b). The chemical structure of PJ1 is 2’-((4-(4-methoxyphenyl)thiazol-2-yl)carbamoyl)-[1,1’-biphenyl]-2-carboxylate (Fig. 2c). The 3D model of PJ1 bound to the OASS pocket (Fig. 2d) shows the interaction of its benzoic acid that is able to compete with SAT I314 followed by an aromatic ring that competes SAT V313 and is linked by amide to an additional aromatic ring that competes with SAT Y312 (Fig. 2d-e).

**Fig 2.**
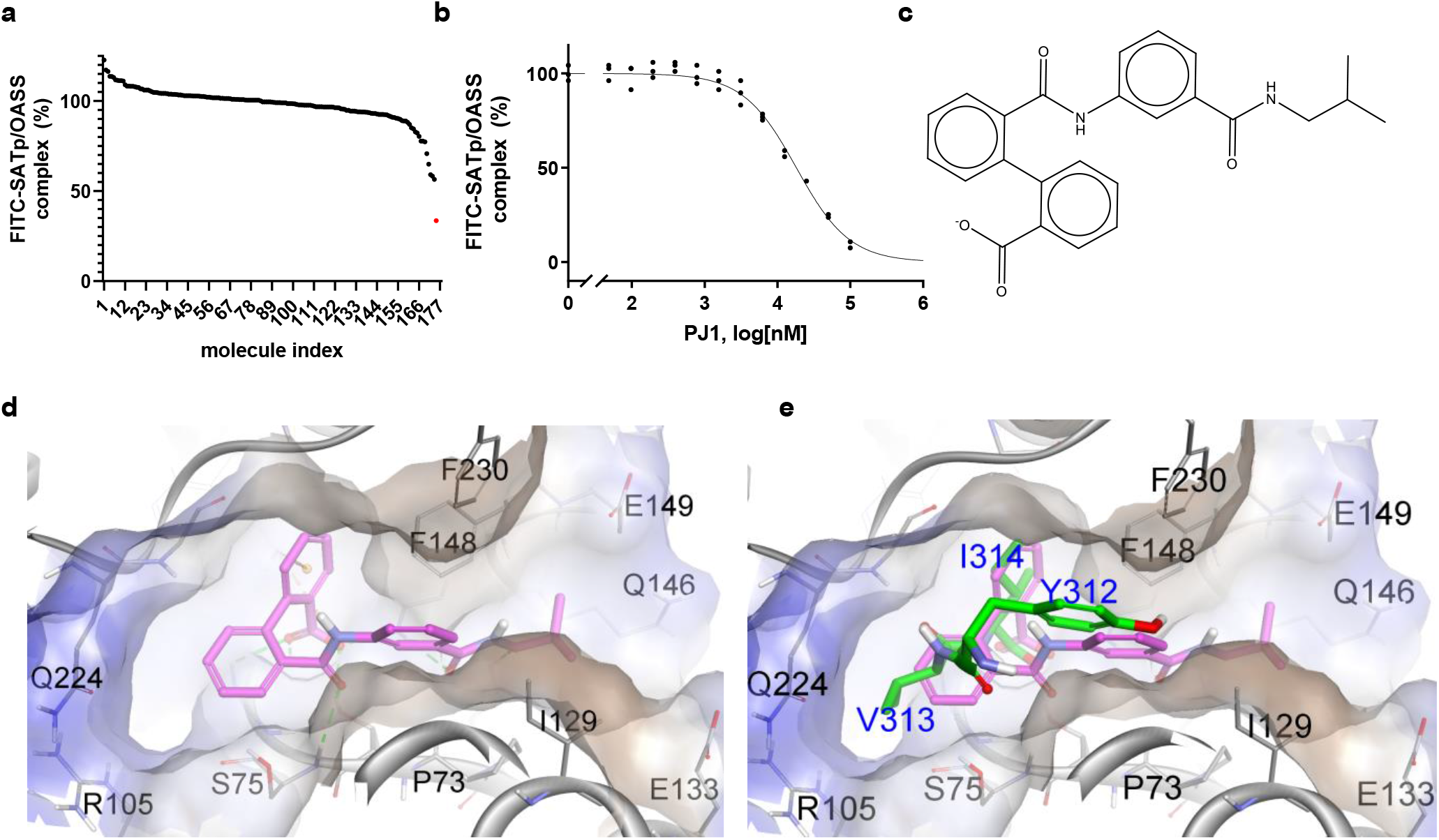
In-vitro screening and characterization of virtual hits. **a**, Evaluation of selected virtual hits by a single 50 μM dose fluorescence polarization (FP) response. mP values were normalized relative to the free FITC-SATp (0%) and bound to OASS with no inhibitor (100%). Data are arranged in descending order of mP value where each hit is labeled by a successive number (x-axis). The most potent hit (PJ1) is color-coded in red. **b**. Full dose-response FP curve of PJ1. Each concentration was measured in triplicates. **c**, Chemical structure of PJ1. **d**, 3D Structural model of OASS with PJ1. **e**, Superposition of SAT (green) amino acids V313, I314 and Y312 with PJ1 (pink), showing the ability of each of the functional moieties in PJ1 to compete with SAT.

### Evaluation of molecules structurally-similar to PJ1

Following dose-response validation, we tested PJ1 analogs to study the functional moieties that best contribute to the binding of OASS and could mark a molecular starting point for further *de-novo* synthesis. As an initial step, we searched for a focused set of available compounds and devised a set of relevant chemical analogs by docking and ranking the molecules using the same principal parameters of the virtual screening. At the end of the process, we purchased ten commercially available molecules containing 2’-carbamoyl-[1,1’-biphenyl]-2-carboxylate, the central part of PJ1 that binds to OASS (PJ2-PJ11, Extended data Fig. 1a). Three of the structurally-similar molecules and their interactions with OASS are shown in Fig. 3a-c.

**Fig. 3.**
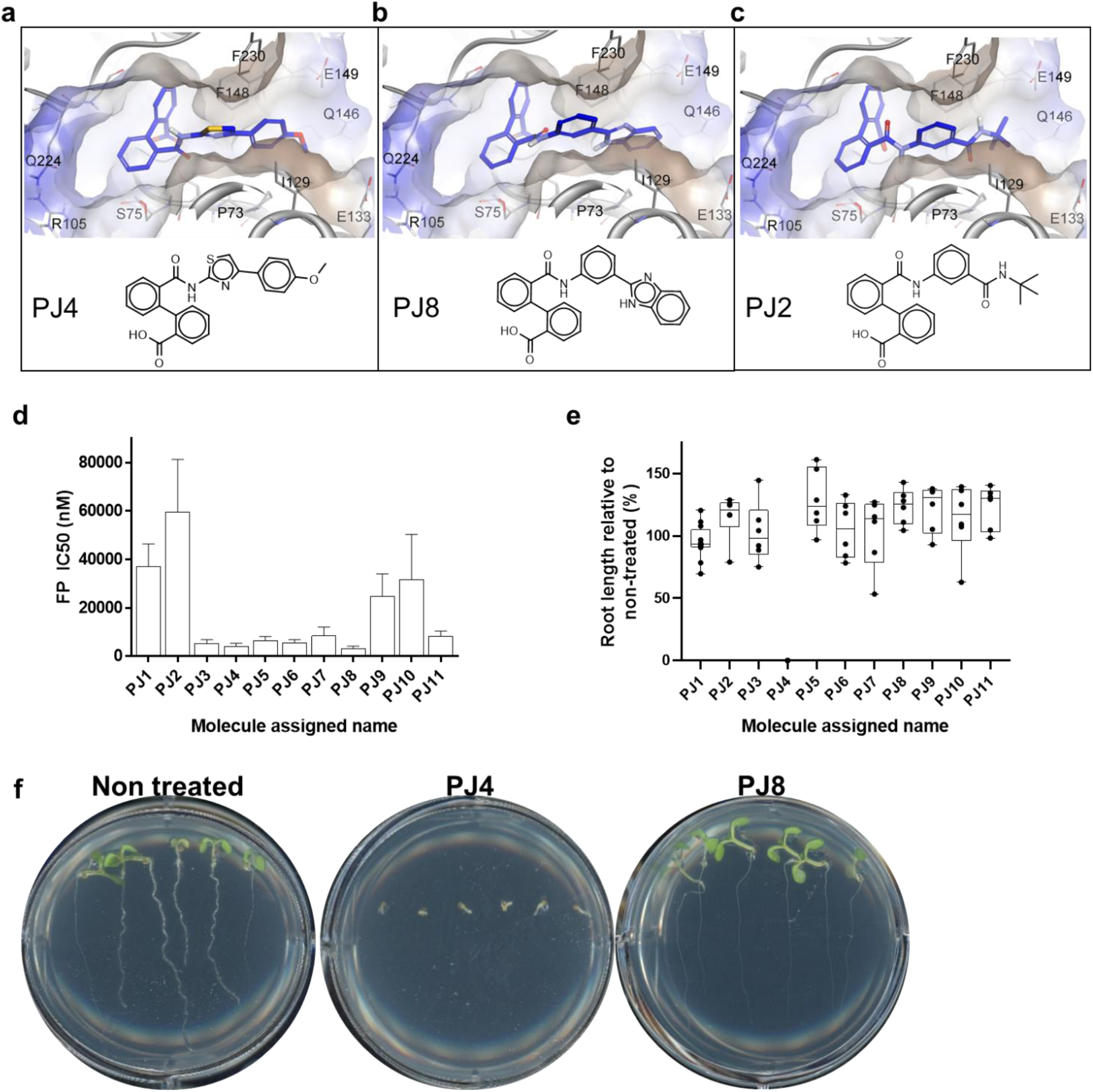
Evaluation of PJ1 structural similar molecules. **a-c**, Representative chemical structures and 3D model with OASS of structurally-similar PJ1 molecules. **d**, IC_50_ values of PJ1-11 elucidated by FP dose-response experiments. **e**, Relative root length of Arabidopsis seeds in agar plates mixed with 50 μM of each compound. Each root is represented by a single dot. **f**, Arabidopsis seedling on non-treated agar plate and agar mixed with 50 μM PJ4 (active) and PJ8 (non-active) after six days. Representative wells taken from different 6-well plates that were tested are shown.

The chemical structure of the compounds reveals several distinctive functional groups, such as methoxyphenyl-thiazole, N-(tert-butyl)benzamide or phenyl-1H-benzo[d]imidazole. To study the effect of each group within the context of the different molecules on the ability to compete with SAT/OASS interaction we tested each molecule in the FP assay. All the molecules competed with the FITC-SATp/OASS complex albeit over a large range of IC_50_ values (Fig. 3d). The FP IC_50_ values of all molecules are summarized in Extended data Table 1 and full dose-response FP curves are shown in Extended data Fig. 2.

### Determining initial in-planta activity via the inhibition of Arabidopsis root growth

To determine which molecules will be the basis for further optimization, we evaluated the *in-planta* activity of structurally-similar molecules (PJ2-PJ11) by measuring their inhibition of elongation of intact roots of germinating Arabidopsis seeds on agar plates. Even though Arabidopsis-based assays do not necessarily correlate with future successful herbicidal activity, they provide a fast and effective evaluation for the initial *in-planta* activity of non-formulated compounds^31^. To this end, Arabidopsis seeds were germinated on agar plates pre-mixed with 50 μM of each molecule and after six days, the root length of each seed was measured and compared to roots in a DMSO-only (vehicle) treated plate. The relative root length of Arabidopsis seeds growing on agar mixed with the different molecules is shown in Fig. 3e. Only PJ4 effectively inhibited Arabidopsis seedlings’ growth (Fig. 3f). The response to 50μM of all tested molecules on Arabidopsis is shown in Extended data Fig. 3. Among this group of analogs able to compete with the FITC-SATp/OASS complex interaction, only PJ4 also inhibited in-planta Arabidopsis seedling growth. Percentage of inhibition by the different molecules are summarized in Extended data Table 1.

### Chemical synthesis of de-novo molecules and structure-activity relationships

Given the improved IC_50_ and in-planta activity of PJ4, we designed and synthesized new chemical entities based on its structural model with OASS. To this end, the four aromatic rings of PJ4 were chemically modified. The synthesized molecules and their modeled binding structure with OASS are shown in Extended data Fig. 1b-d. Representative molecules are shown in Fig. 4a. First, the substitution of the ring with the methoxy group of PJ4 (Fig. 2a) that occupies the OASS secondary pocket distant to the SAT peptide (Fig. 1a) was explored. Although the secondary distant pocket is hydrophilic, PJ4 and PJ1 contain methoxy and isobutyl groups, respectively. We thus aimed to evaluate the affinity of molecules with a hydrophobic group. To this end, we substituted the PJ4 methoxy group with hydrophobic groups (e.g., PJ14, PJ15). However, the substitution of the methoxy to di-chloro (PJ15) or to alkyl (PJ14) groups did not improve the FP IC_50_ (IC_50_=4.7 μM and 5.4 μM, respectively). Adding a chlorine in the ortho position together with methoxy in the para position (PJ19) did improve binding affinity resulting in IC_50_=3.6 μM. In addition, substitution to positively charged groups was explored (e.g., PJ13, PJ12 and PJ16). Such a charge can potentially interact with OASS E133, E149, or E72. Indeed, the substitution of the methoxy with an amino ethyl (PJ13), (phenoxyethyl)morpholine (PJ12), or nitrophenyl (PJ16) resulted with IC_50_= 1.9, 1.9 and 2.6 μM, respectively.

**Fig. 4.**
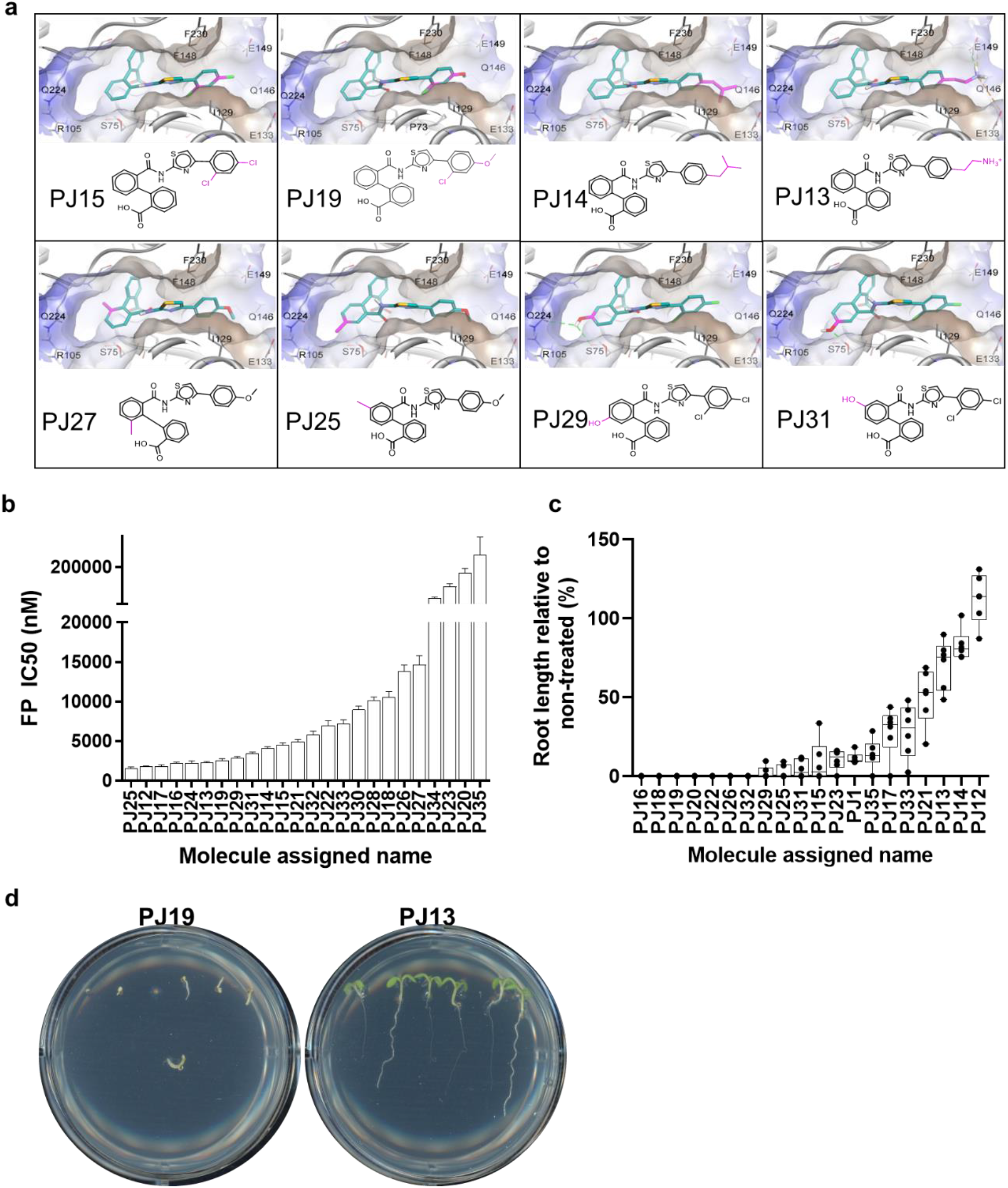
Arabidopsis growth inhibition. **a**, Designed molecules based on PJ4 chemical structure. **b**, IC_50_ values of the de-novo synthesized molecules were evaluated in FP experiment. The bars represent the IC_50_ value (nM) calculated for each molecule. **c**, Relative root length of Arabidopsis seeds in agar plates with each of the indicated molecules in a concentration of 50 μM. Each root length is represented by a single dot. **d**, Two representative images of Arabidopsis seedlings on agar plates treated with PJ19 (active) and PJ13 (non-active) are shown. Representative wells taken from different 6-well plates of the same experiment are shown.

The second functional region of PJ4 that was explored for substitutions is the ring that competes with SAT V313. Analysis of OASS pocket in which V313 is located shows it can accommodate a larger group than the current aromatic ring in PJ1 and PJ4. Several alternatives were explored to better occupy this pocket, such as addition of a methyl group at the ortho (PJ27), meta (PJ24) or para (PJ25) positions on the carboxylic ring. While the limited volume substitution at the ortho position seem to have failed disrupt the OASS/SAT interaction (IC_50_=18.5 μM), the substitutions at the meta and para positions were slightly better (IC_50_=3.4 and 2.1 μM, respectively). As the pocket occupied by V313 is hydrophilic, we also sought to increase the number of hydrogen bonds with S75, R105 or Q224. To this end, we added hydrogen donor/acceptor hydrophilic groups to the ring to compete with SAT V313, such as hydroxy and amide groups. Indeed, PJ31 hydroxy in the para position interacts with the S75 side chain leading to IC_50_=4.5 *μ* M and PJ29 with a hydroxy group at the meta position interacts with the S75 backbone and the Q224 side chain leading to IC_50_=3.4 μM. This improvement that shows greater binding affinity relative to PJ15 affirms the importance of such chemical alterations. Acetamide or hydroxymethyl groups at the meta position (PJ30 and PJ28, respectively) are probably too bulky, leading to an IC_50_=11 and 9.3 μM, respectively. We also tried to replace one carbon in this ring with nitrogen (PJ18 and PJ26) or by replacing the ring to oxazol (PJ20 and PJ35), hypothesizing it may serve as an acceptor of hydrogen bond with S75. However, these substitutions resulted in weaker binding affinities. The third functional group of PJ4 that we explored is the benzoic acid that competes with SAT I314. Two molecules were synthesized with methoxy in para position relative to the carboxyl (PJ21) or with fluorine in the meta position (PJ22). Both interact with OASS but with nearly no effect on IC_50_ relative to PJ4. The fourth functional group that was explored is the thiazole ring. Methyl substitution of the thiazole (PJ33) did not improve the competition with SATp. In addition, more drastic changes like the alteration of the thiazole in PJ4 with (1H-pyrazol-1-yl)ethan-1-ol (PJ34) or pyridine-2(1H)-one (PJ23) hampered the ability of the molecules to compete with FITC-SATp/OASS, resulting with an IC_50_=50 and 34 ***μ***M, respectively.

These preliminary structure-activity relationship studies indicate that the core structure of our molecules interferes with the binding of the SAT C-terminus that is anchored within OASS binding site. The benzoic acid of the molecules competes with SAT I314. However, the substitution of this ring did not improve binding affinity. The benzoic acid followed by an additional 6-membered aromatic ring competes with SAT V313 and cannot be replaced by a 5-membered ring. However, a 6-memebred ring can be substituted bearing small hydrophobic or hydrophilic groups. The thiazole group that competes with SAT Y312 should be linked to one more aromatic ring. These two planar rings impart the molecules with the ability to penetrate and fill the region that is not occupied by SAT. The latter ring can be substituted by a hydrophobic group like the methoxy of PJ4, or better with a positive group like PJ13.

### In-planta activity of second-generation de-novo synthesized compounds

To further filter the newly designed molecules and evaluate *in-planta* activity, we repeated the single-dose Arabidopsis root growth inhibition assay with the newly synthesized molecules (Fig. 4c). Although some of which, such as PJ12 and PJ13, were able to compete with the FITC-SATp/OASS interaction, they did not have significant in-planta activity. Other molecules such as PJ20 did not effectively compete with the complex formation but yielded a significant in-planta activity, probably due to off-target effects. Such differences between direct binding and cellular activity often arise due to poor cell penetration, variable solubility, or fast chemical alteration of the molecules within the cell. Representative images of Arabidopsis roots treated with two indicated molecules PJ19 and PJ13 are shown in Fig. 4d. The images for the single-dose Arabidopsis root inhibition of all molecules are shown in Extended data Fig. 3.

Molecules that were active in the single-dose experiment were further evaluated to elucidate the *in-planta* IC_50_. Representative images of seedlings on agar plates with variable concentrations of PJ4 and the corresponding root length in each concentration are shown in Fig. 5. The in-planta IC_50_ of all tested compounds is shown in Extended data Fig. 4 and data are summarized in Extended data Table 1. As often occurs following a single-dose screening, some of the molecules did not show repeatable activity results (e.g., PJ29).

**Fig. 5.**
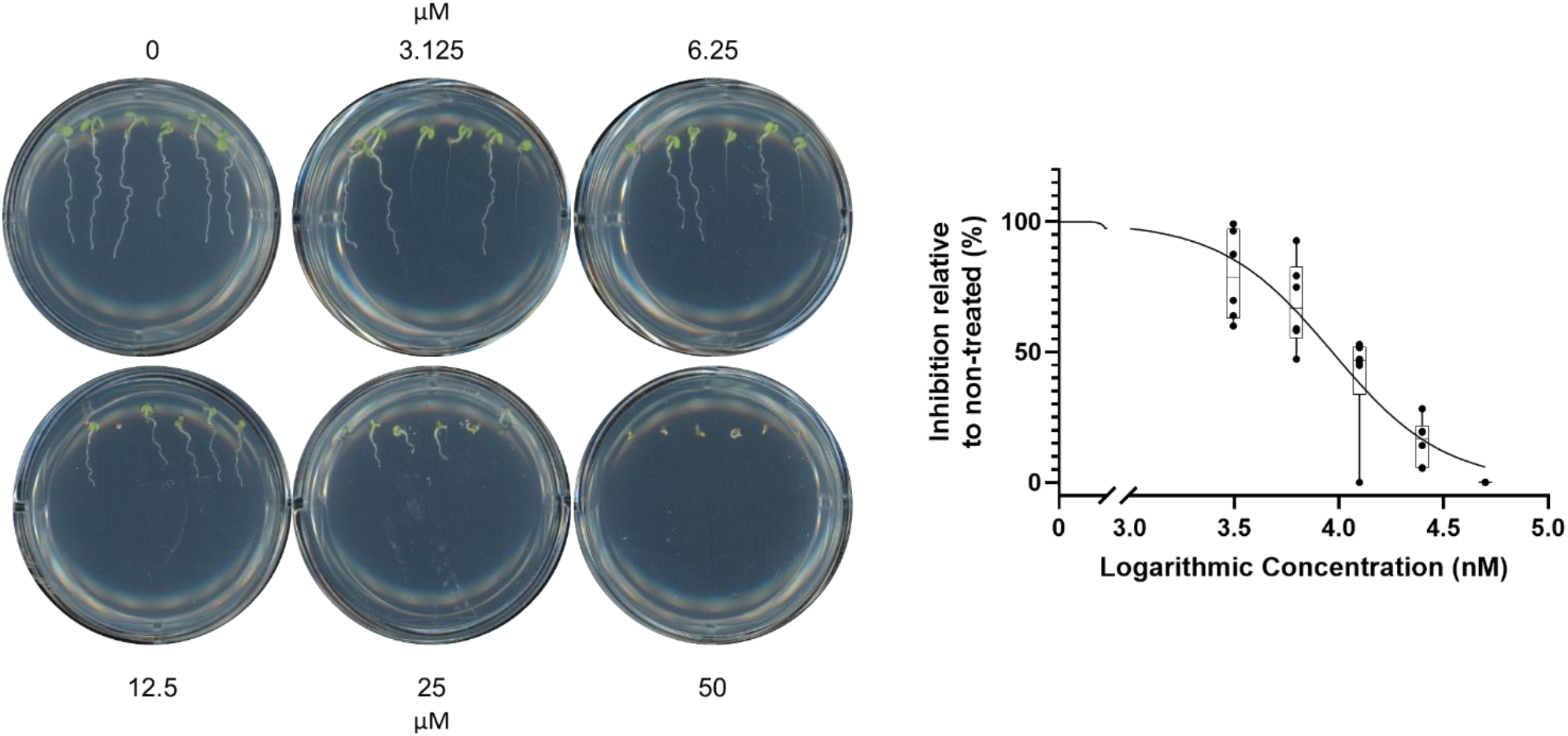
Images of Arabidopsis seedlings treated with variable concentrations of PJ4 and elucidation of the IC50. Representative wells taken from different 6-well plates in the same experiment are shown.

### In-planta pre-emergence activity evaluation

To further study the in-planta activity of active and inactive molecules, we evaluated the molecules’ ability to inhibit the emergence of Clover and Amaranthus. Since such applications, especially of the non-formulated active ingredient, often require higher concentrations of the molecules, the latter were synthesized in their salt form to allow their dissolution to a final concentration of 5 mM in H_2_O. Following seed planting, the soil was sprayed with 2 ml yielding an approximal final concentration of 4.6 kg/H of the non-formulated compounds. Representative images of the Clover pots 21 days after spraying each molecule (right pot) and vehicle (left pot) onto the soil are shown in Fig. 6. Oxyfluorfen, a commercially available herbicide, was sprayed as a positive control at a concentration of 5 mM, corresponding to 3.6 kg/H, according to the manufacturer’s recommendations. Extended Data Fig. 5a-b show the pre-emergence experiments on Clover and Amaranthus, respectively, growing on soil treated with various molecules.

**Fig. 6.**
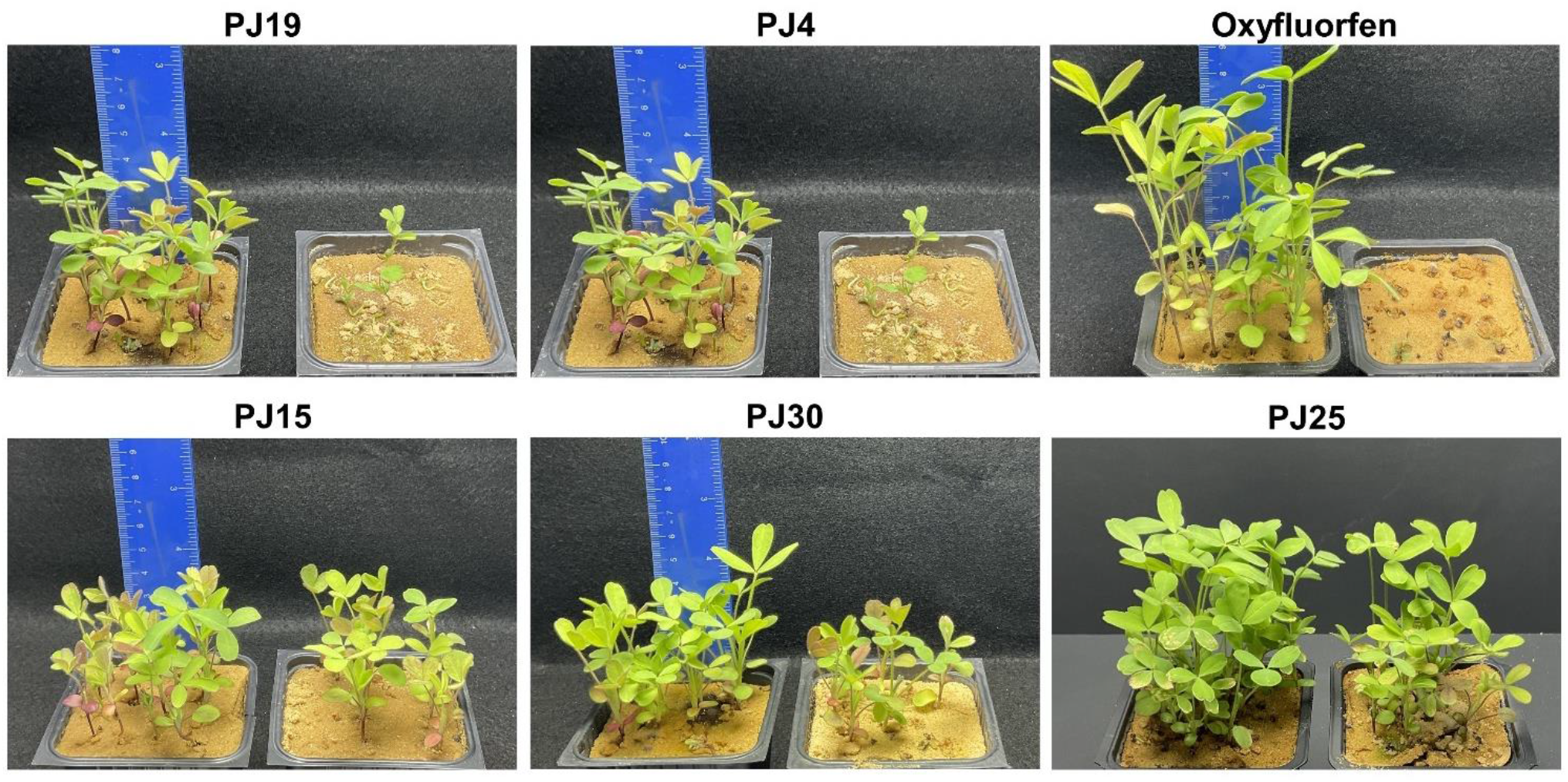
Pre-emergence inhibition of Clover by representative chemical leads. Molecules were sprayed onto the soil following seed planting. Pictures were taken 21 days after planting. Chemical structures of these chemicals are shown in Extended data Fig. 1.

### Evidence that the lead chemical PJ4 inhibits cysteine biosynthesis

To further validate the mechanism at the cellular level, we tested the ability of PJ4 to inhibit cysteine biosynthesis in cultured tobacco BY-2 cells. The sequence and structure of the OASS/SAT interaction surface is highly conserved among plant species, it is expected that inhibitors targeting this epitope will also be active in the tobacco cells. BY-2 cells were cultured with and without 25 μM of PJ4, and the levels of free cysteine were measured by HPLC. Free cysteine levels were reduced by 70% (Extended data Fig. 6) as expected.

## Conclusions

Our results demonstrate that it is possible in a short time to determine hot spots in protein-protein interactions, rapidly screen a virtual library of existing and potential chemicals, evaluate their in-vitro protein binding activity and actually find phytotoxic active lead pesticidal molecules. Some leads, even when not optimally formulated showed in planta activity and can be the basis of further chemical synthesis and formulation to increase activity. This computational-based pesticide discovery was done with a minimum of syntheses and orders of magnitude of less greenhouse space compared to the conventional ‘spray-and-pray’ discovery and screening that has led to no commercialized new herbicide mode of action in decades.

The OASS-SAT interaction interface exemplifies an important plant PPI. Three main reasons pose PPI as attractive herbicidal targets. The first is that the number of PPIs is much higher than that of single proteins, presenting a huge number of yet unexplored protein interfaces that have important cellular roles. The second is that a weed would have to evolve simultaneous complementary mutations in the hot spot of both partners for the enzyme complex to retain activity while repelling the chemical. Such an event renders the evolution of weed target site resistance in the field highly unlikely. Indeed, our preliminary modeling shows that the majority of amino acid substitutions in the SAT binding site that preclude the binding of our chemical lead compound (PJ4), would result in the SAT not interacting with its OASS partner (Extended data Fig. 7). A limited number of mutations indicated in the inset could still hamper the binding to PJ4 while still maintaining binding to OASS. Further chemical optimization is thus needed to eliminate any potential mutation. Whereas such simultaneous mutations are unlikely in the field, it is possible to model what mutations would retain enzymatic activity while preventing the binding of the chemical. This could assist for the development of transgenic crops resistant to the chemical. An additional motivation of PPI relates to vertebrate biosafety. The third motivation is that within the realm of PPI, those involved in the synthesis of amino acids, and in particular the essential ones for mammals, are an attractive target for the development of biosafe herbicides. Indeed, we previously showed that molecules inhibiting the methionine biosynthesis pathway are inhibitory in-planta^11^.

Altogether, our study succeeded in elucidating small molecules that bind OASS, enabling the inhibition of OASS-SAT enzyme activity and plant growth. We used a multidisciplinary approach in which the integration of computational and biophysical/biochemical tools enabled us to screen a large library of small molecule candidates and validate the activity of the selected virtual hits toward OASS. This is of immense importance when targeting PPI interfaces that were considered “undruggable”. Together with these achievements, some open questions still remain. We have not demonstrated that OASS is the sole target of our lead chemicals, this can further be shown by having plants with OASS dual mutants that are active in the presence of our lead chemicals. This is also important for studying off-target activity and how it will affect mammals. In addition, questions remain related to the practical application of the molecules in the field, large-scale synthesis cost of the molecules, their stability and environmental effects.

Our study indicates the potential ease and efficiency of successful development of pesticidal lead compounds having new modes of action, by targeting a protein-protein interaction (PPIs). Our findings demonstrate the potential for rapid identification of active molecules and provide a foundation for further chemical syntheses to enhance activity. Such methodology could pave the way to overcoming many of the safety and resistance challenges that are associated with current pesticides.

## Methods

### Profiling and filtering of small molecule virtual library

Library profiling was based on pharmacophore constraints based on the analysis of the interaction between SAT C-terminal and known peptide and small molecule binders to OASS based on protein data bank (PDBs): 2ISQ, 3ZEI, 3IQH, 3IQG, 3IQI, 2Q3B, 2Q3D, 2Q3C, 1Y7L, 1D6S, 4LMB, 5XOQ, 5J43, 5J5V, 4ZU6, 4ZU1, 4LI3, 5DBH, 5DBE, 4ORE, 4NU8, 3T4P, 3TBH, 4JBN, 4IL5. In addition, following classification and analysis of molecular features of known herbicides. Pharmacophore hypothesis and filtering were done by the search DB algorithm in Catalyst program^32^. Small molecules that fulfilled all required features within the geometrical constraints of the pharmacophore model were then ranked by “fitting value”—a calculated score based on the distance between all features of the model, and chemical group position of each small molecule. The result of this step was a reduction of the library size to approximately 400K compounds.

### Molecular Docking and Final Compound Selection

OASS structure (PDB ID:2ISQ) preparation and forcefield assignment was carried out using CHARMM ^33^, and molecular docking of the profiled library was done using the discovery studio software, with preset genetic algorithm settings optimized for virtual screening. A post-docking scoring function was applied to rank each docking pose. A cutoff value was used to filter out compounds that did not satisfy specific location constraints and also to account for putative ligand interacting groups in addition to the structure-based complementary ones.

### OASS expression and purification

The DNA of *Arabidopsis thaliana* OASS (uniprot P47998) was synthesized and optimized for *E. coli* expression (Genscript ltd.). The gene was cloned with an inducible promoter into a pGex-6p-1 plasmid containing an N-terminus GST (glutathione transferase) tag followed by a PreScission Protease cleaving site. The vector was transformed to *E. coli* Rosetta cells and cultured to OD=0.8. Protein expression was induced by 0.5 mM IPTG and the cells were cultured for an additional 16 h at 25°C and harvested. Following cellular lysis, the soluble fraction was passed through a GST column and the bound GST-tagged protein was eluted by 20 mM reduced glutathione. The GST tag was cleaved by overnight dialysis incubation with PreScission Protease and the mixture passed again through the GST column to separate the OASS from the GST and PreScission Protease constructs. OASS purification and molecular weight were validated on SDS-PAGE gel.

### Fluorescence polarization

Fluorescence measurements were made on samples arrayed in 96-well plates using an F-200 Tecan® polarizer. To screen small molecules for their ability to compete with OASS/SAT, a FITC-labeled SAT peptide (YLTEWSDYVI) was synthesized. The peptide was incubated with 2 μM OASS in the presence of each evaluated molecule and following 30 min of incubation fluorescence was read at an excitation/emission of 485/535 nm. The raw data were fitted to a single site inhibition model by Graphpad Prism 7. All measurements were executed in triplicates.

### BY-2 cell culture

Tobacco BY-2 cells were cultured in sterile Murashige and Skoog basal medium (Sigma-Aldrich, cat#: M0404), supplemented with 0.2 μg/L 2,4-dichlorophenoxyacetic acid, 100 μg/ml myo-inositol, 1 μg/L thiamine HCl and 3 % sucrose. The cells were incubated in the dark at 25 °C on a shaker at 130 rpm. Cells were diluted (1:15) in fresh media every seven days. Before treatment with each molecule, the cell suspension was adjusted to OD(600 nm)=0.4-0.5 and 100 ml seeded in erlenmeyer flasks. PJ4 was added at a final concentration of 25 μM and cells were cultured for 24 h.

### Determination of cysteine levels by HPLC

The cells were filtered on Whatman No. 1 paper to remove excess liquid and then lyophilized for 24 hours to complete dryness. Twenty milligrams of treated and non-treated BY-2 cells were ground into a fine powder using liquid nitrogen, and disrupted in a Restch MM 400 homogenizer. The powder was suspended in 400 μl of 0.1 M HCl to precipitate proteins followed by two cycles of centrifugation for 10 min at 4 °C at 14,800x*g*. The resulting supernatants was stored at -80 °C until cysteine content detection by reverse-phase high-performance liquid chromatography (HPLC). 25 μl from each sample was neutralized with 25 μl of 0.1 M NaOH and 1 μl of 0.1 M dithiothreitol (DTT) was added for disulfide bond reduction. Derivatization of the sulfide group was carried out with 50 μl of monobromobimane thiol-reactive fluorescent probe (MBB) solution (1 mM MBB in 200 mM Tris-HCl buffer pH 8). The solution was incubated at 37 °C for 15 min in the dark. The MBB labeling was stopped with a final concentration of 4.5% acetic acid. After centrifugation for 4 min at 4 °C at max speed the samples were filtered through PVDF 0.2 μm filter. Separation and quantification of thiols were carried out on a Kromasil 100 C18, 250 × 4.6 mm column using the UHPLC (UltiMate 3000, Dionex) system. HPLC run was based on a two buffers gradient with 0.1 % acetic acid (buffer A) and 100 % acetonitrile (buffer B) as follows: i) 10 to 15% B for 10 min, ii) 15 to 20% B for 5 min, iii) 20 to 100% B for 2 min, iv) 100% B for 7 min and, v) 100 to 10% B for 1 min. The bromobimane thiol derivatives were detected fluorometrically (ex/em 390/482 nm). L-cysteine (SIGMA) was used as a reference standard and treated in the same manner as the sample supernatants. Peaks at approximately 4.8 min were identified as cysteine. Data is the average of three biological repeats, each performed in replicates.

### Inhibition of *A. thaliana* seedlings

Seeds were washed with 70% ethanol for 1 min and then with sterile water. Surface sterilization was achieved using a solution containing 2.5% sodium hypochlorite and 0.2% Triton X-100 for 10 min followed by washing five times with sterilized water. Half strength Murashige and Skoog medium (MS; Duchefa, Haarlem, Netherlands), supplemented with 0.8% agar and 1% sucrose (w/v), pH 5.8 in a single or six-well plates were prepared. Selected small molecules were dissolved in the agar mixture to a final concentration of 50 μM with a remaining 0.1% DMSO in each plate. Then, *A. thaliana* seeds were placed in each well. Plates with the *A. thaliana* seeds were placed at 4 °C for 48 h and then transferred to a growth chamber for 6-7 days (22 ± 2°C) at day/night cycles of 16/8 h. The plates were stored vertically. The control well contains 0.1% DMSO. For the dose–response assay, molecules were diluted to the desired concentration and transferred to the medium. At the end of incubation time, plates were scanned, and root length was measured using ImageJ software. The percentage of inhibition was calculated by averaging all root lengths in a plate.

### Pre-emergence in planta experiment

*Amaranthus biltum* (16 seeds each) or *Trifolium alexandrinum* (clover; 25 seeds each) were planted in pots filled with 20% garden soil (bottom layer) and 80% sand (upper layer) covered with sieved dry sand. The spray solution containing 5 mM of each molecule in 10% acetone, were sprayed in a chemical hood. The time of spraying was dependent on the germination time of each species; Clover pots were sprayed 1 day after sowing and *Amaranthus* pots were sprayed 3 days after sowing. The pots were placed in a growth room equipped with two LED lamps emitting 200 μEin at plant height at a temperature of 23 °C and day/night cycles of 16/8 h. Plants were irrigated every 2 days for 21 days.

## Author contributions

I.B. and E.C. designed the computational model, performed the computational screening and docking, analyzed the computational data, and selected the virtual hits. R.B.S and N.D. set up the biophysical, cellular *in vitro* assay and in-planta evaluation, and executed all experiments and protein purification assays. N.D, D.P and J.G supported experimental planning and strategy. N.D and J.G wrote the manuscript and gave critical comments. N.D, I.B. and M.G wrote the manuscript, conceived the original research and supervised the research.

## Data availability statement

The authors declare that the data supporting the findings of this study are available within the paper and its Supplementary Information files. Should any raw data files be needed in another format they are available from the corresponding author upon reasonable request.

