## Supplementary Information for "Pesticide chemical leads inhibiting protein-protein interactions"

[#] Equal first authors

**Corresponding Authors**
Maayan Gal;
Itai Bloch;

**Extended Data, Figures caption**

**Extended Data Fig 1**. **Chemical structure and modeling**. Extended data Fig. 1a shows the structural-similar molecules to PJ1 that were explored for the binding of OASS and tested *in-planta* activity to inhibit Arabidopsis root growth. Extended data Fig. 1b-d show de-novo synthesized molecules based on the structure of PJ4. Chemical variations from PJ4 are shown in pink. Extended Data Table 1 summarizes the molecules' SMILES.

**Extended Data Fig 2**. **FP curves**. The panels show the FP curves of molecules measured by competing for the binding of OASS with the FITC-labeled SAT peptide. Extended Data Table 1 summarizes the IC50 values evaluated by fitting the data into a single-site inhibition model. Further experimental details are found in the main text.

**Extended Data Fig 3. Single-dose Arabidopsis root growth assay**. The panels show the effect of molecules tested in a concentration of 50μM for inhibiting Arabidopsis root growth. Further experimental details are found in the main text.

**Extended Data Fig 4. Dose-dependent Arabidopsis root growth assay.** The panels show the effect of variable concentrations of molecules tested for inhibiting Arabidopsis root growth. In-planta $IC_{50}$ values are summarized in Extended Data Table 1. Further experimental details are found in the main text.

**Extended Data Fig 5. Pre-emergence assay.** The panels show the pre-emergence of clover and Amaranthus treated with selected molecules. Further experimental details are found in the main text.

**Extended Data Fig 6. Cysteine levels in BY-2 cells.** The panel shows the relative cysteine levels of BY-2 cell treated with PJ4. Further experimental details are found in the main text.

**Extended Data Fig. 7. Mutational analysis of PASS for the binding of SAT and PJ4**. The figure shows the $\Delta\Delta G$ for the binding energy of OASS with PJ4 (Y-axis) or SAT-peptide (X-axis) following the substitution of each amino acid in the OASS binding region to the different 19 amino acids. Each point represents the energy difference of a single mutation. Red box - possible mutations that destabilize the binding of OASS with PJ4, but stabilize or non-affecting biding to SAT. Stabilizing $< -0.5$, Destabilizing $> 0.5$, Neutral $> -0.5$ and $< 0.5$.
