## Supplementary figures and images for "Pesticide chemical leads inhibiting protein-protein interactions"

### Figure S1

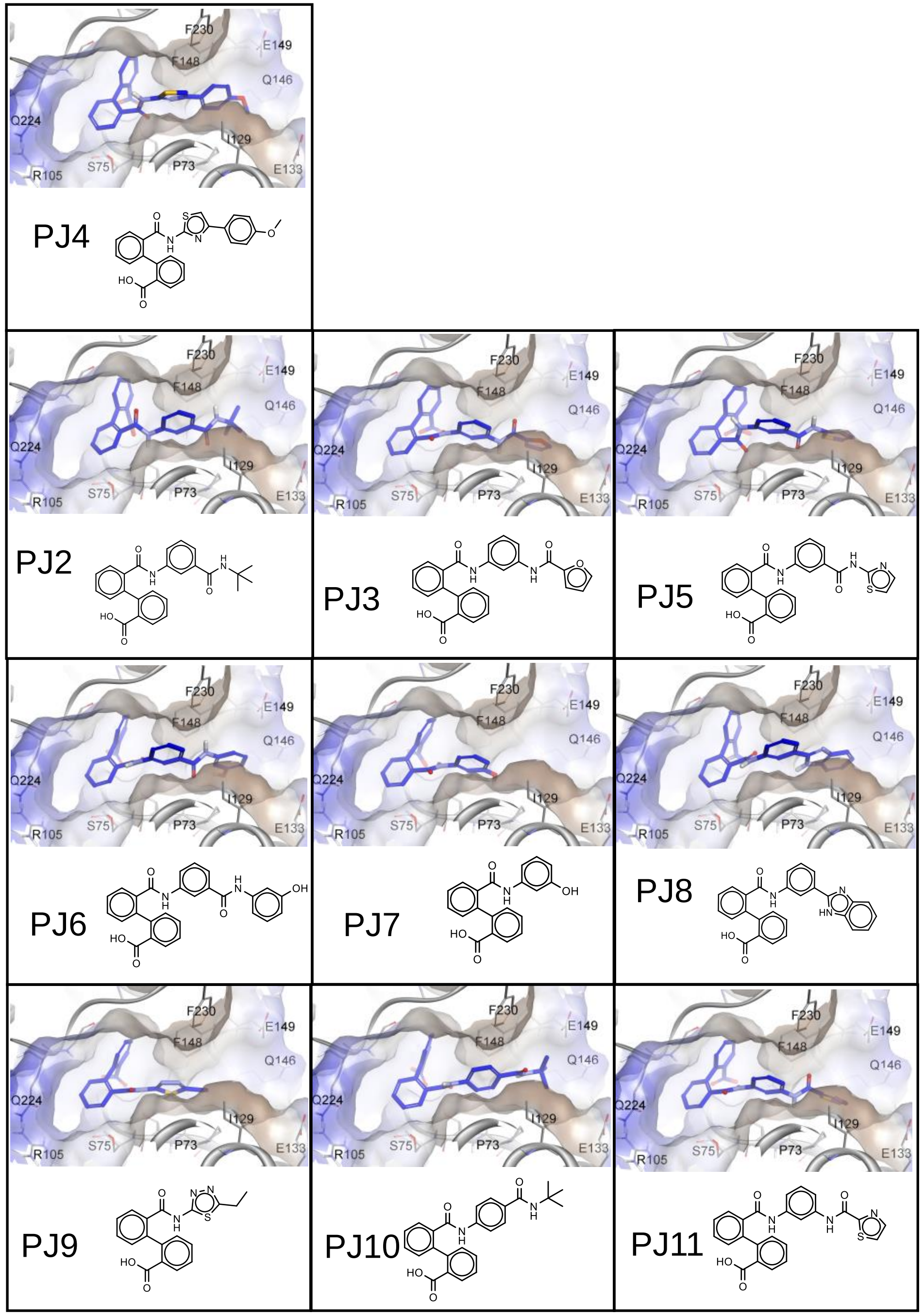


**Extended Data, Fig 1a**

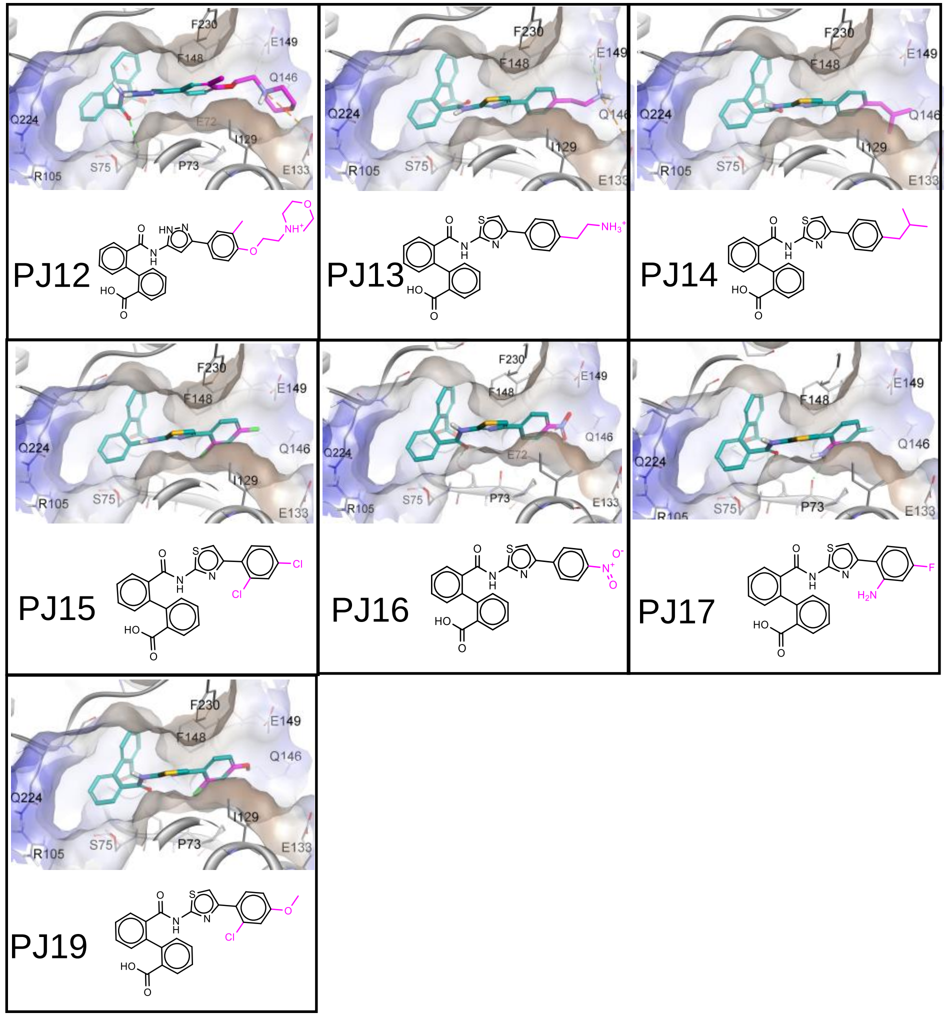


**Extended Data, Fig 1b**

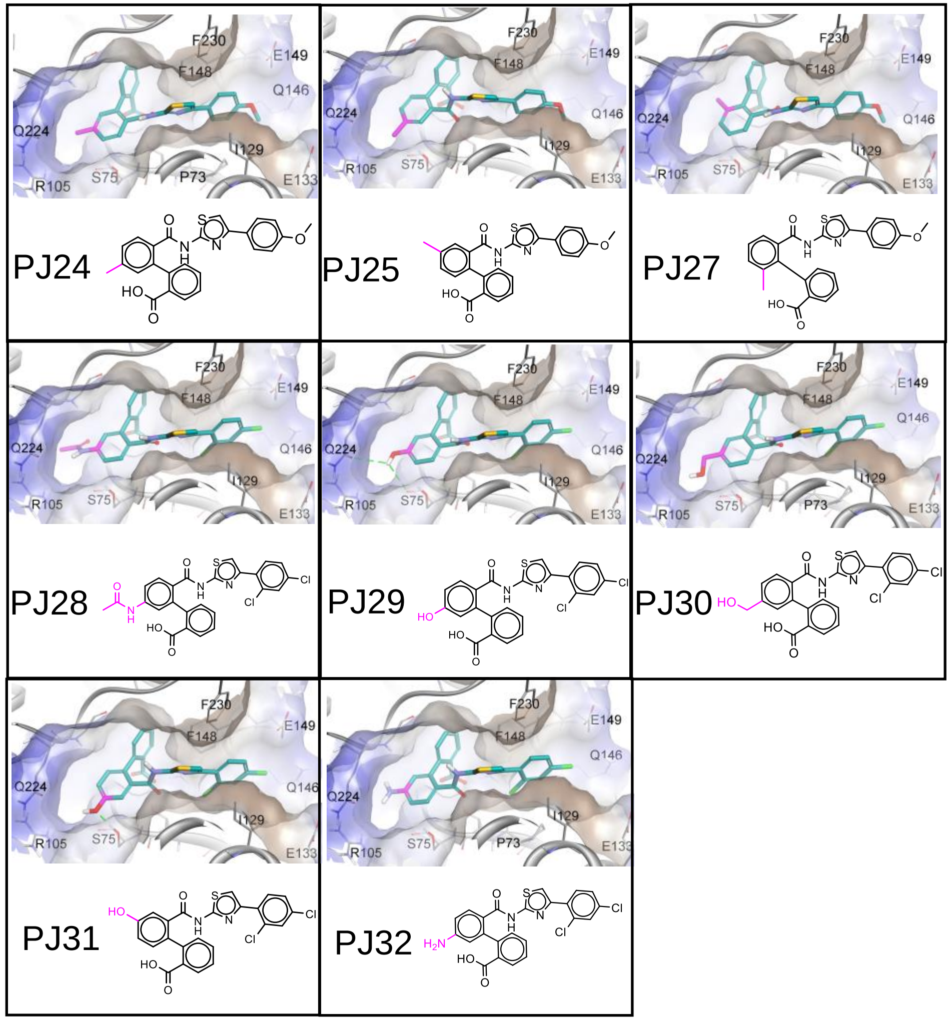


**Extended Data, Fig 1c**

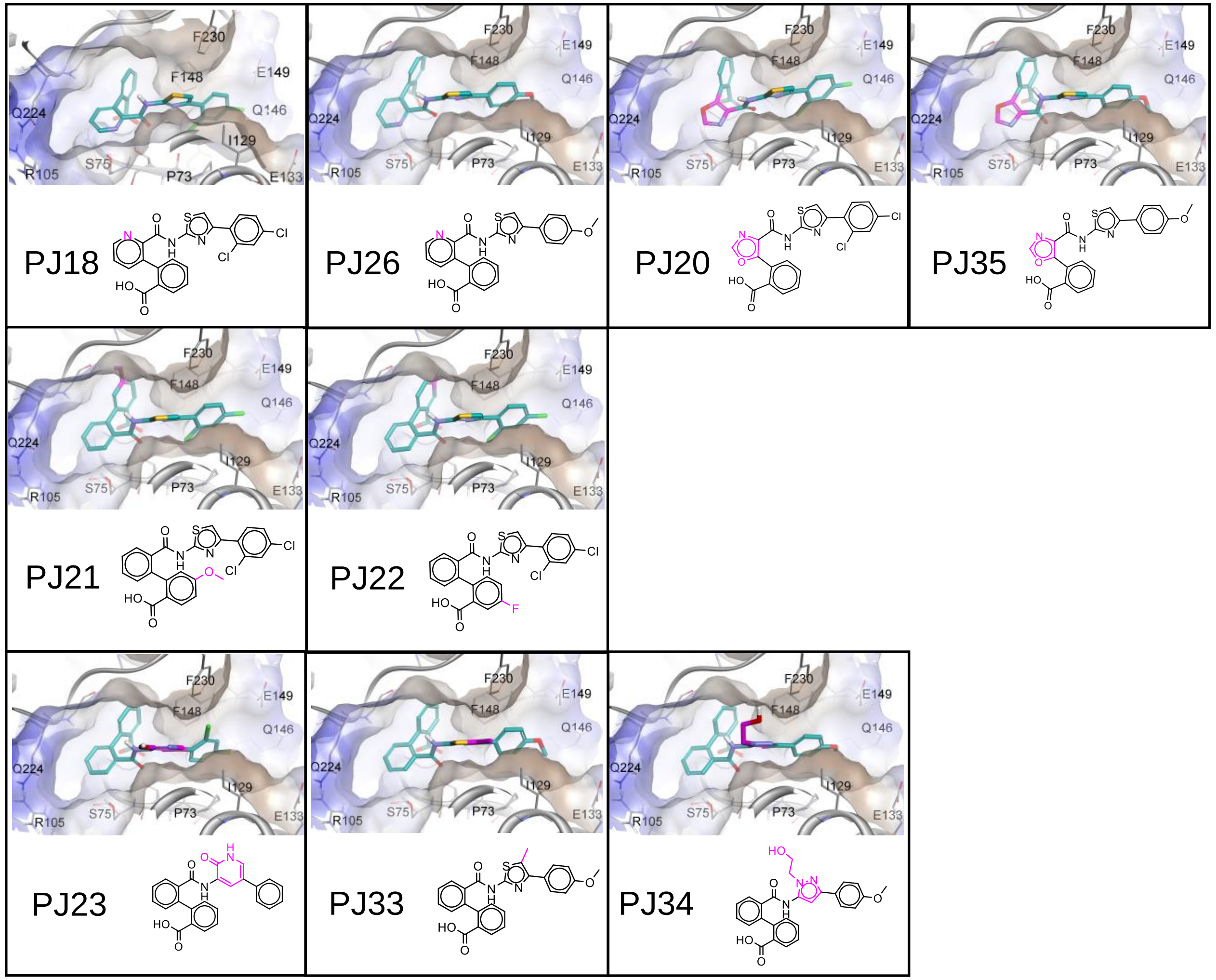


**Extended Data, Fig 1d**

### Figure S2

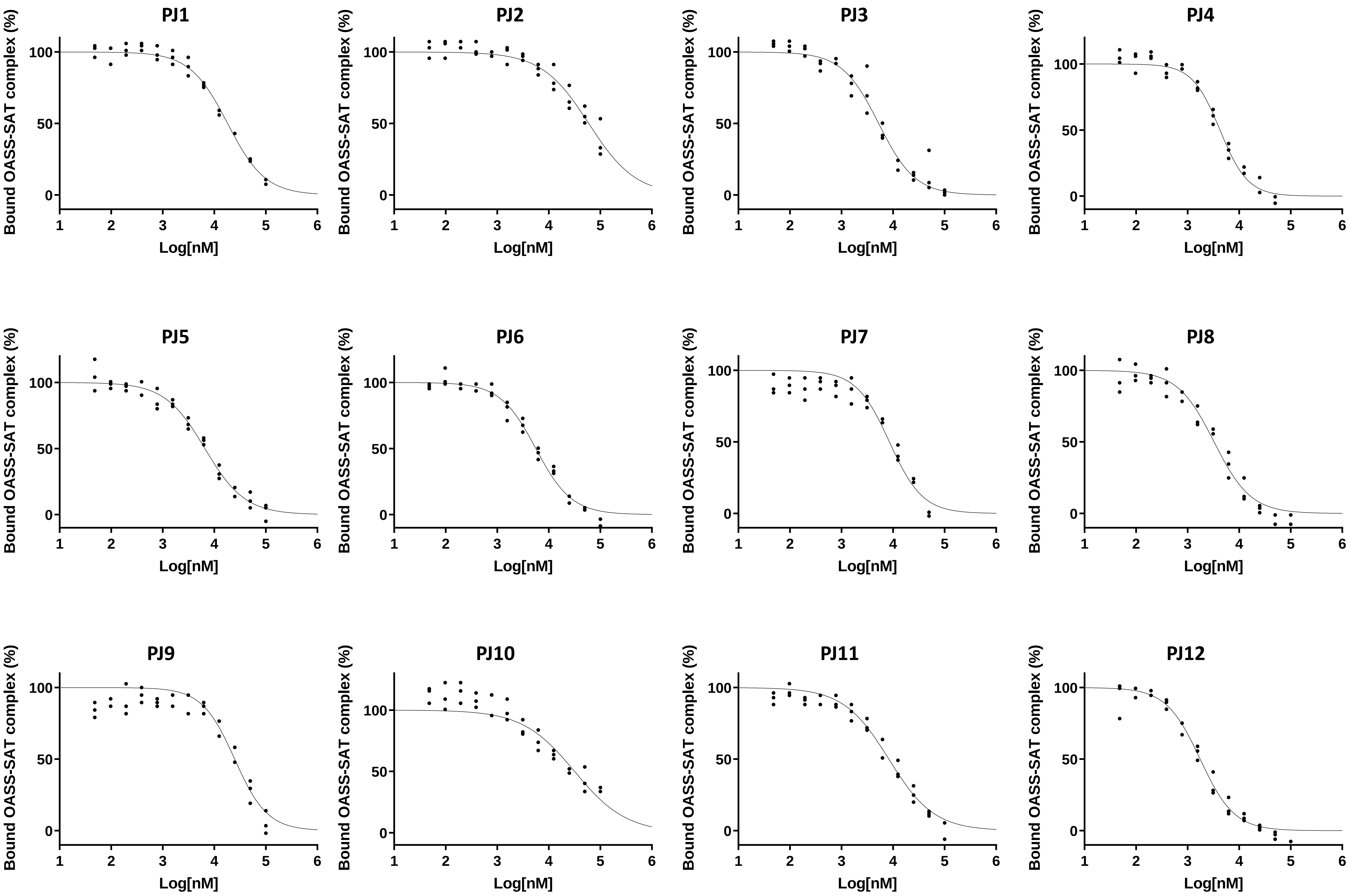


**Extended Data Fig. 2**

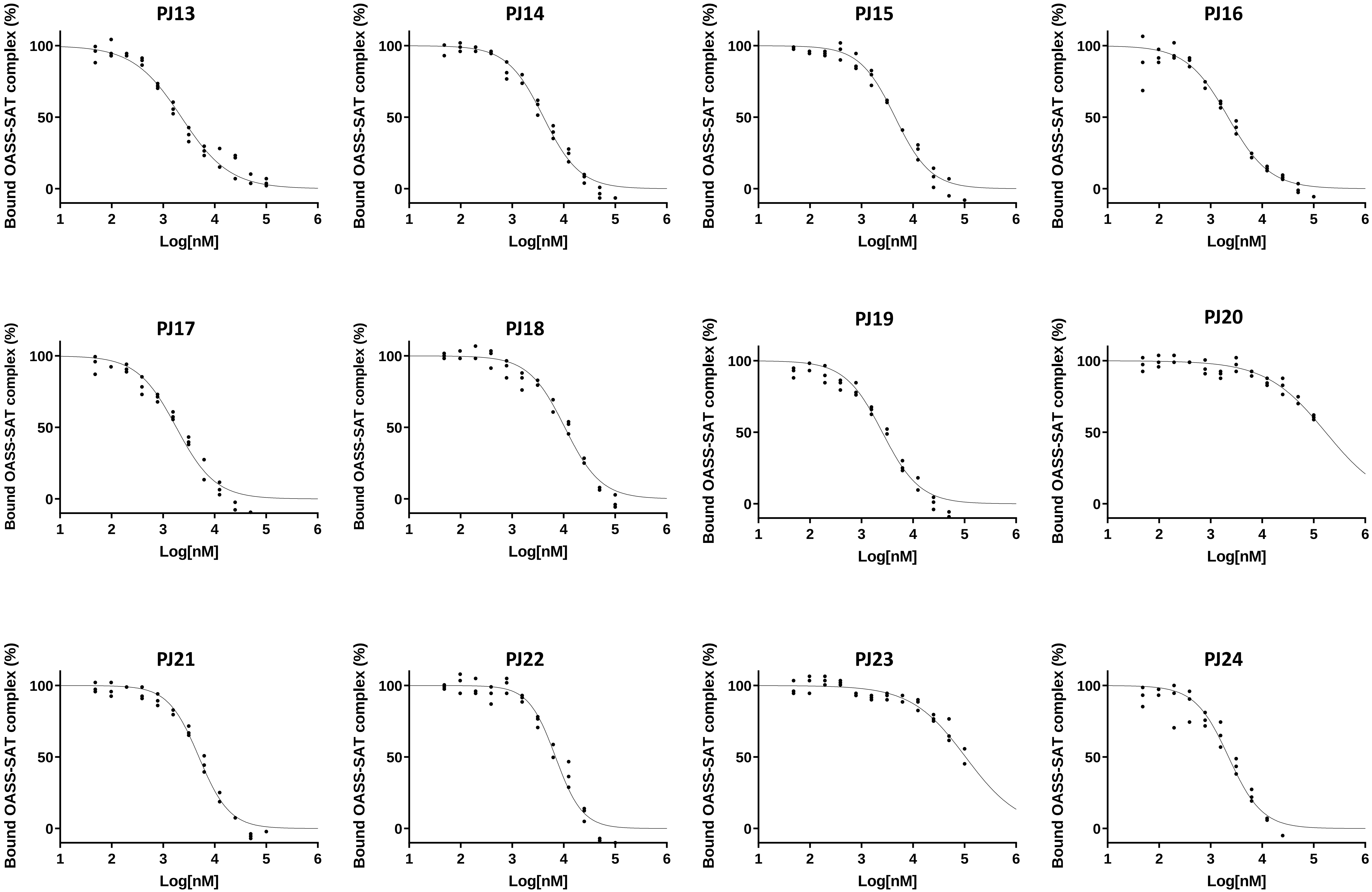


Extended Data Fig. 2

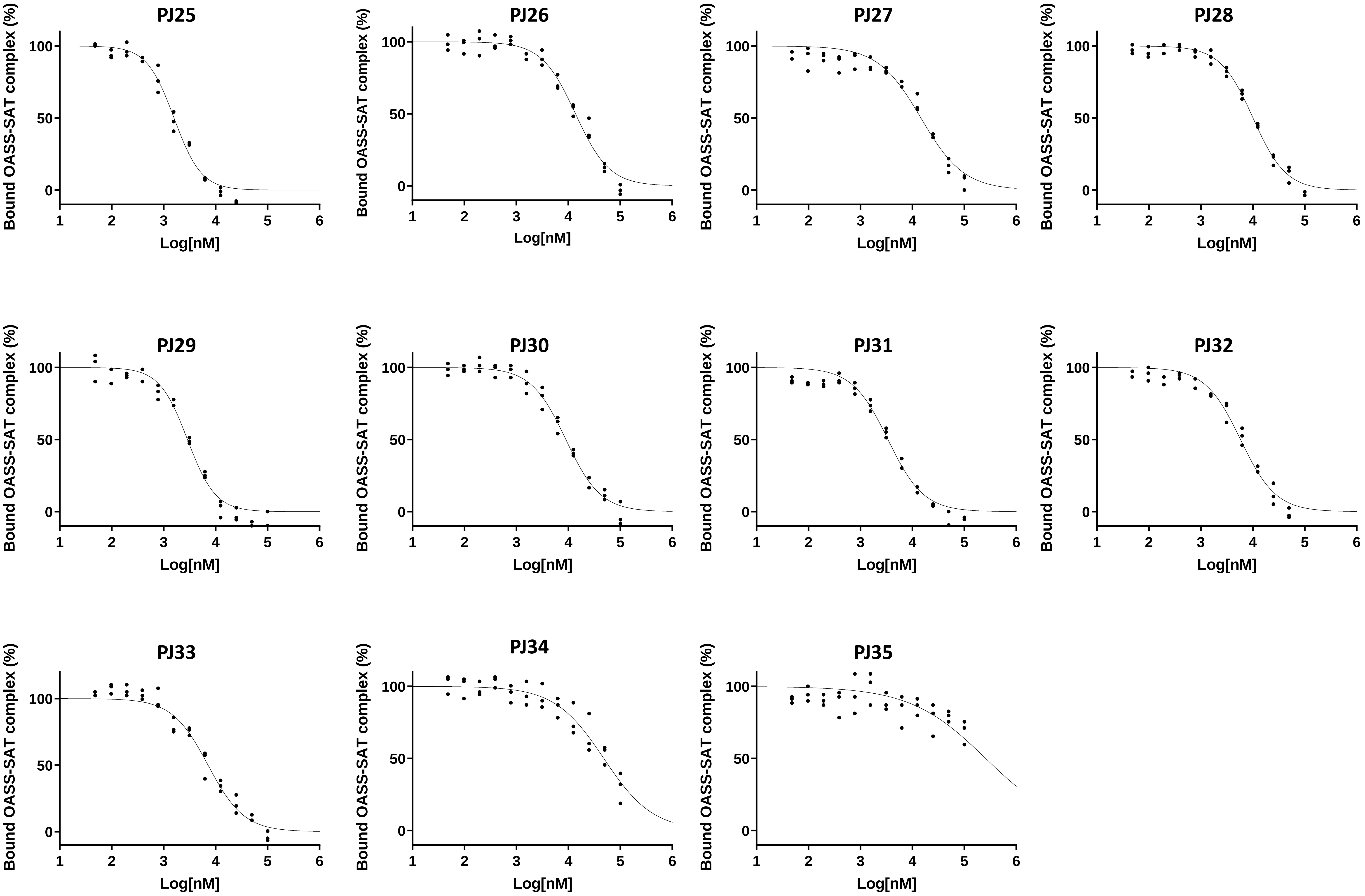


**Extended Data Fig. 2**

### Figure S3

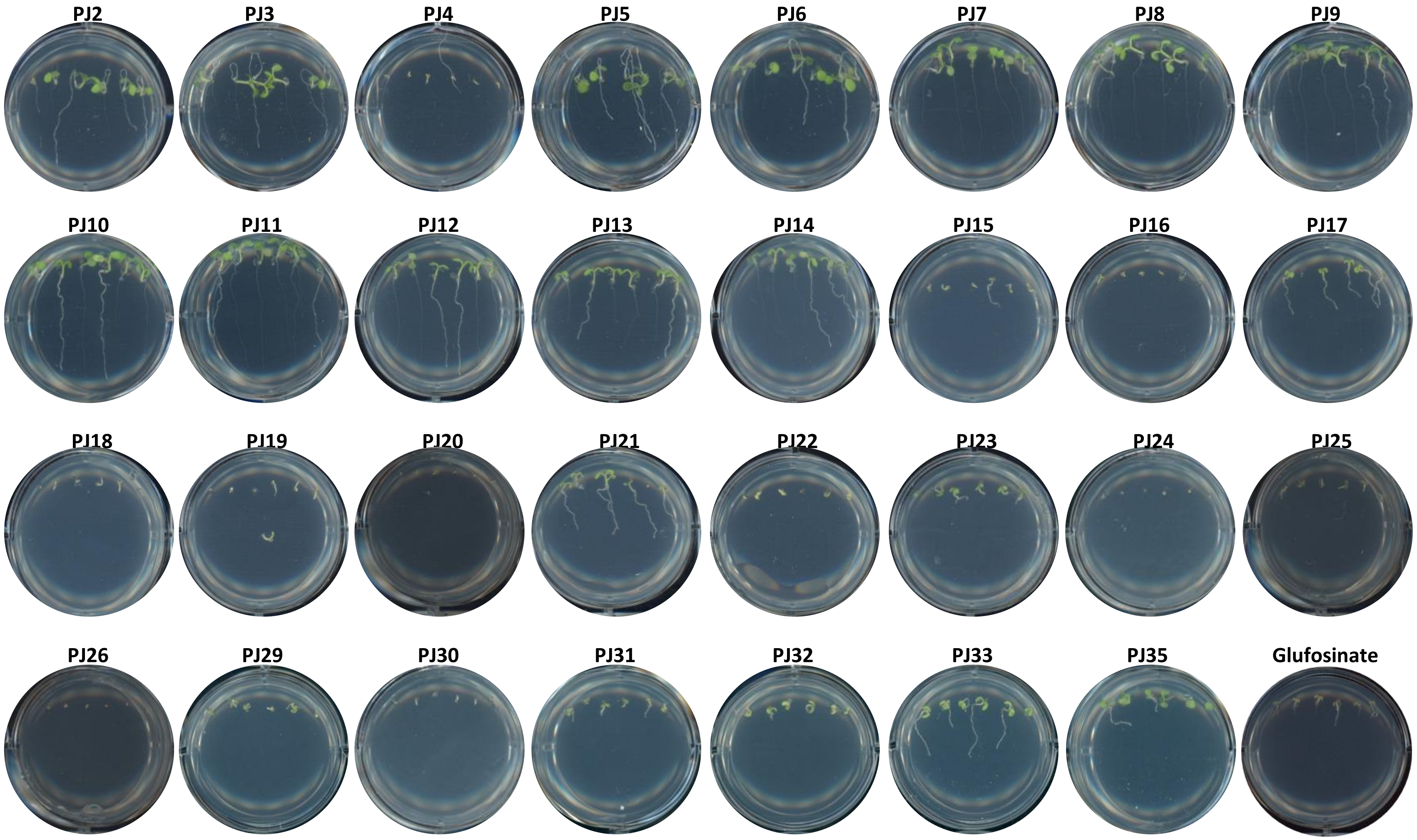


**Extended Data, Fig. 3**

### Figure S5

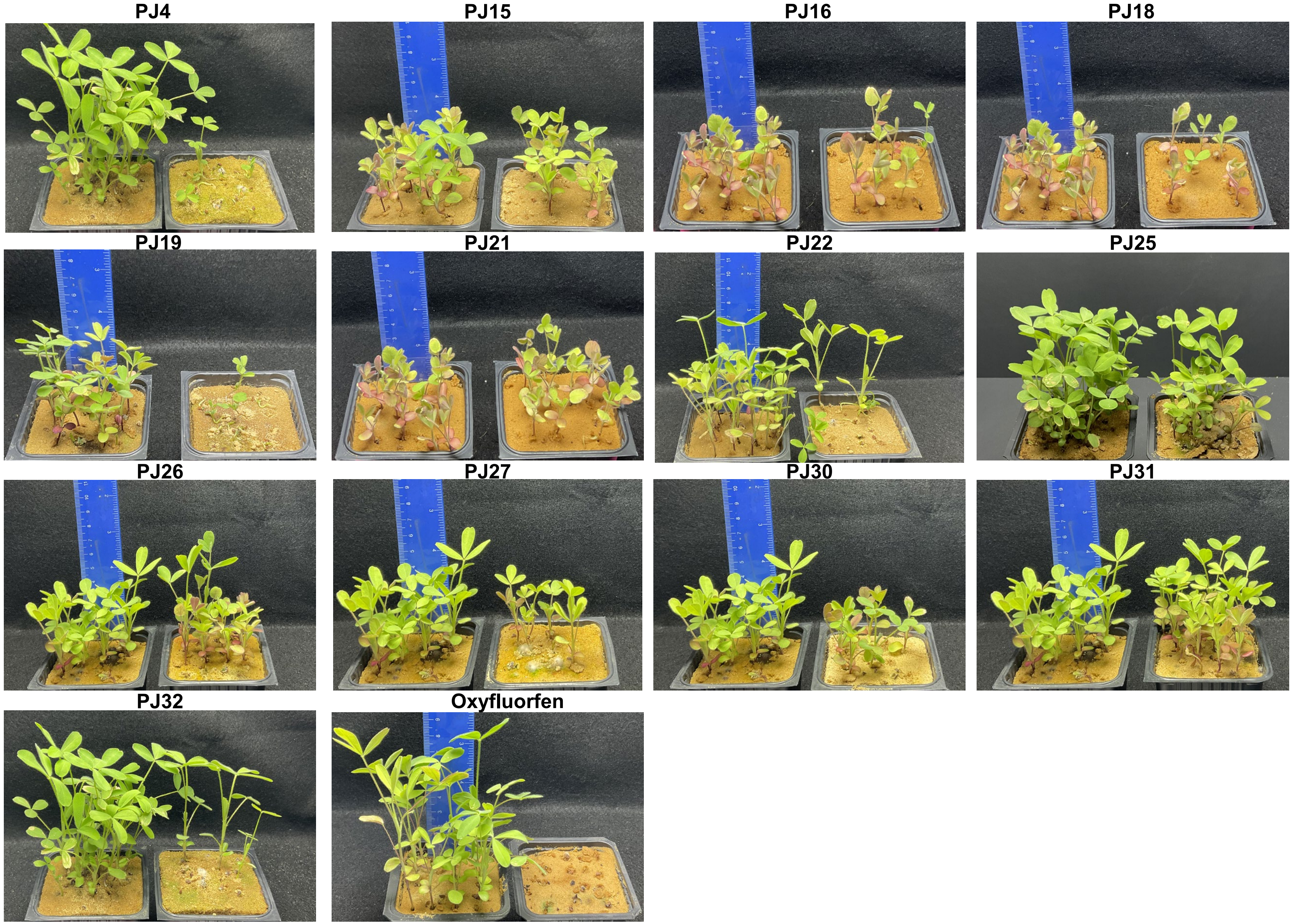


**Extended Data, Fig. 5a**

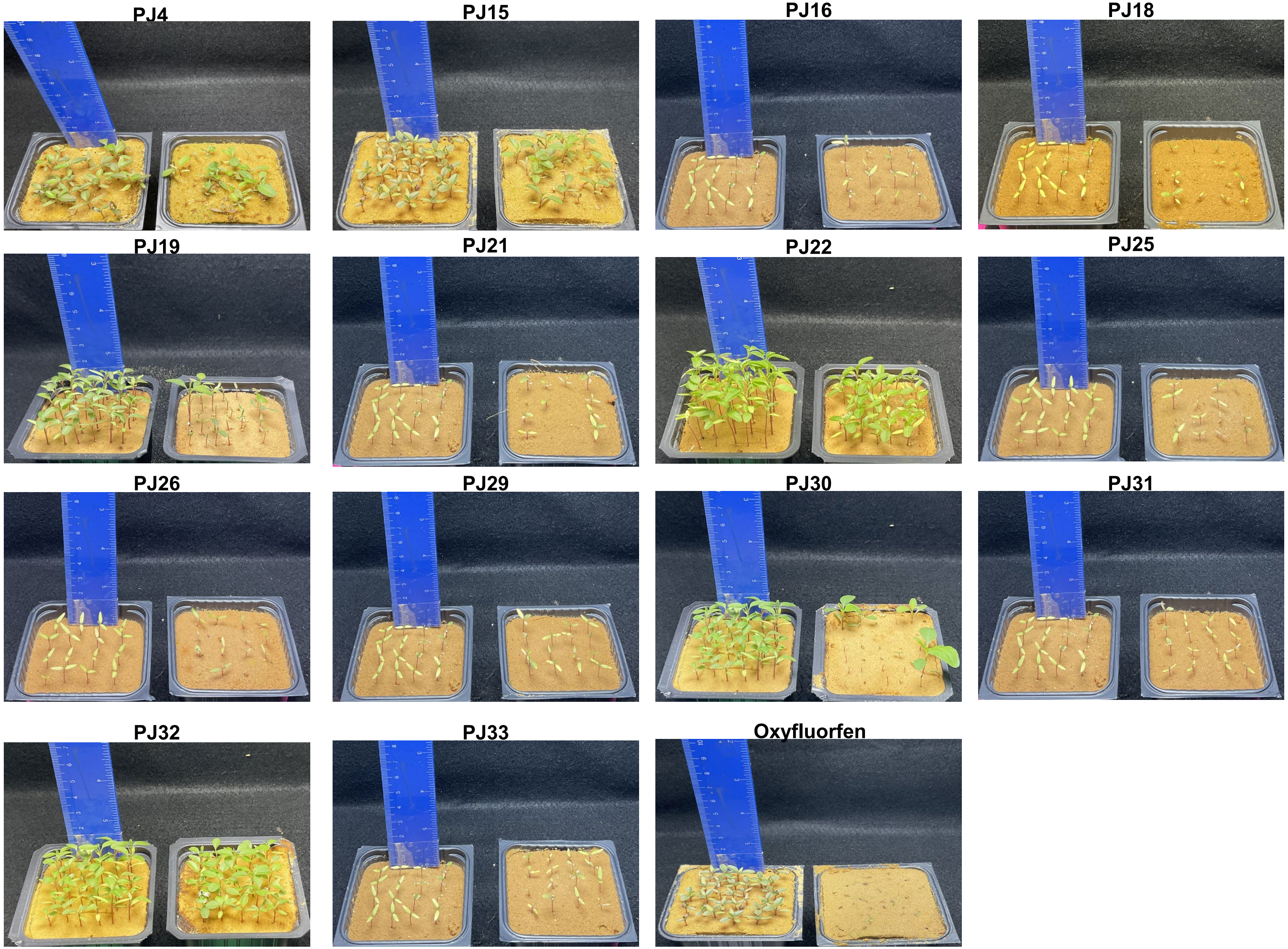


**Extended Data, Fig. 5b**

### Figure S6

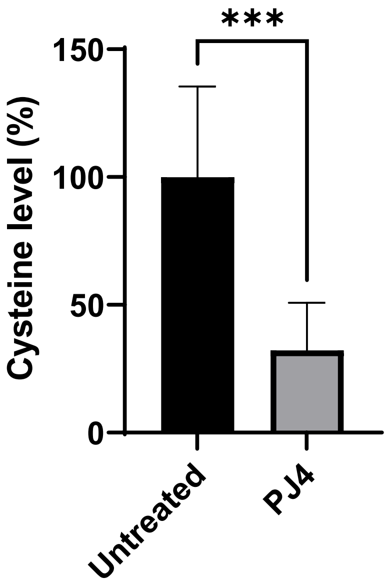


Extended Data, Fig. 6

### Figure S7

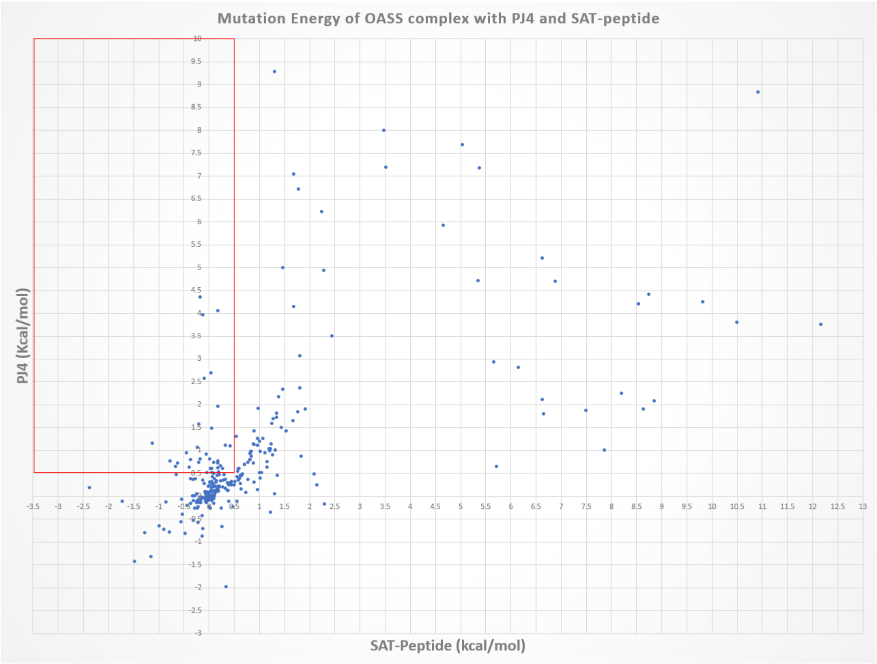


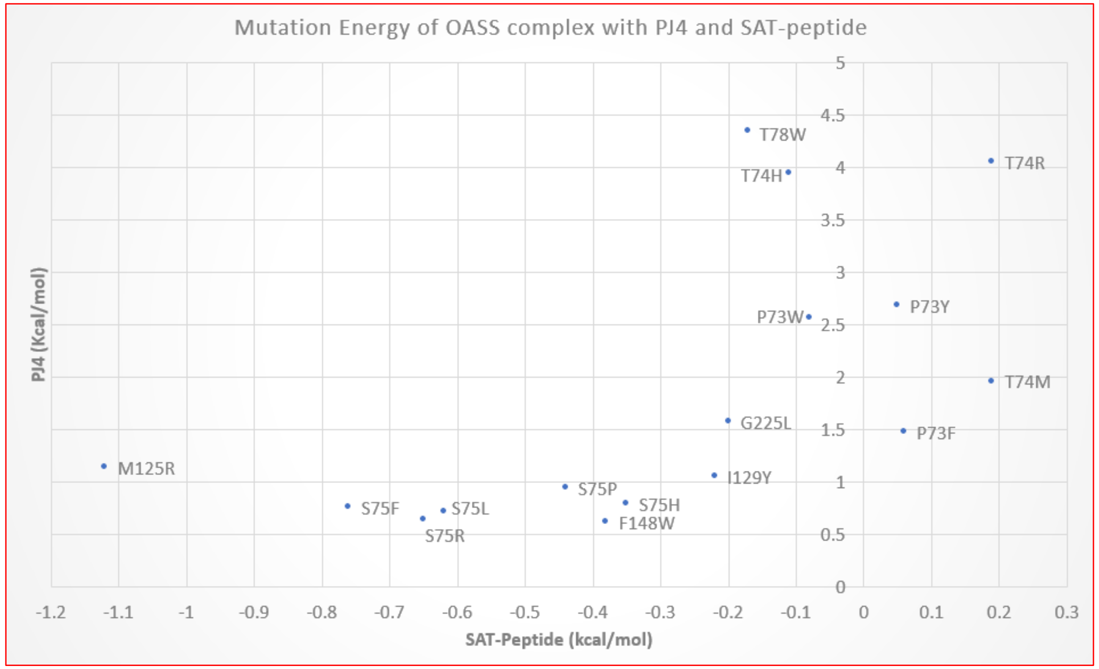


**Extended Data, Fig. 7**
