## Supplementary material for "Pesticide chemical leads inhibiting protein-protein interactions": Figure S4

PJ4

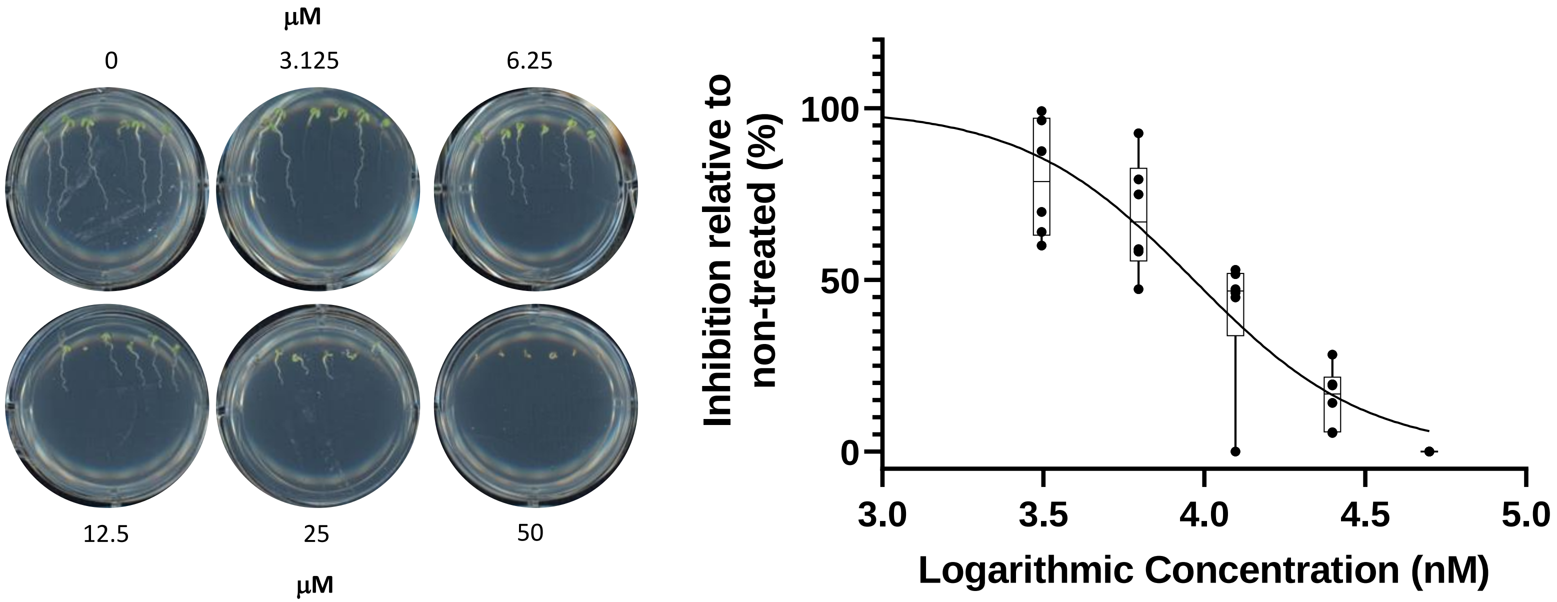


**Extended Data Fig. 4**

PJ14

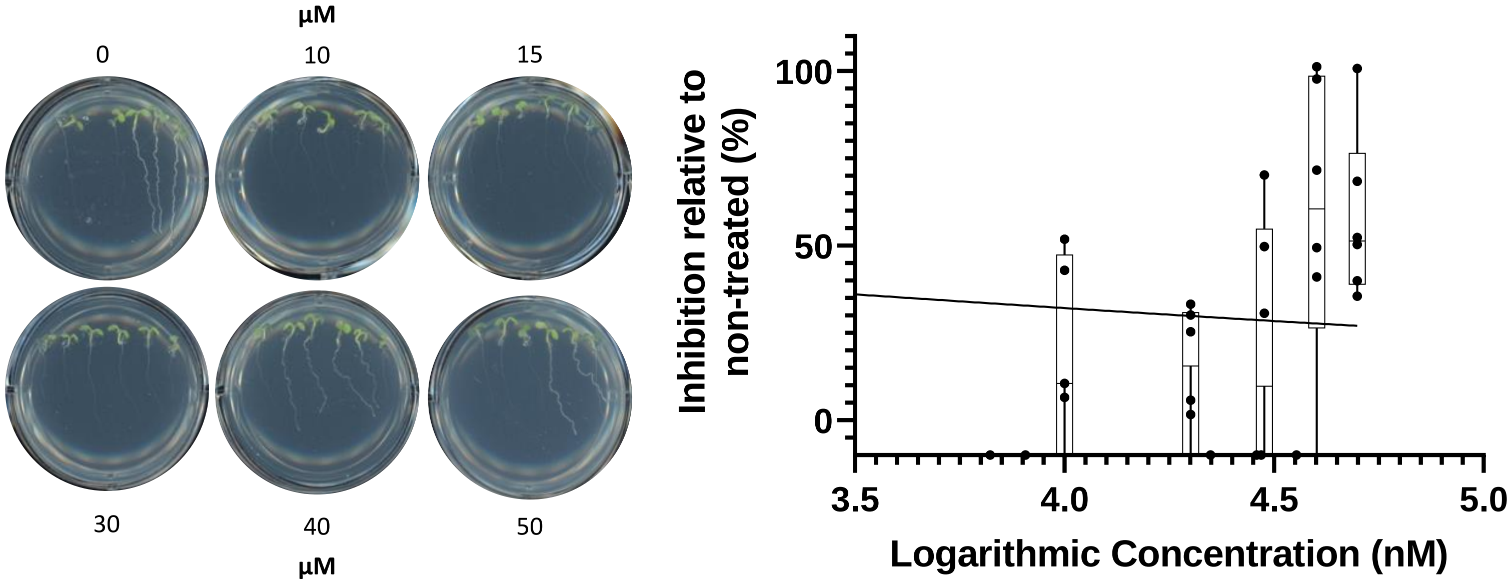
μM
0
10
15
30
40
50
μM
Inhibition relative to non-treated (%)
100
50
0
3.5
4.0
4.5
5.0
Logarithmic Concentration (nM)

**PJ15**

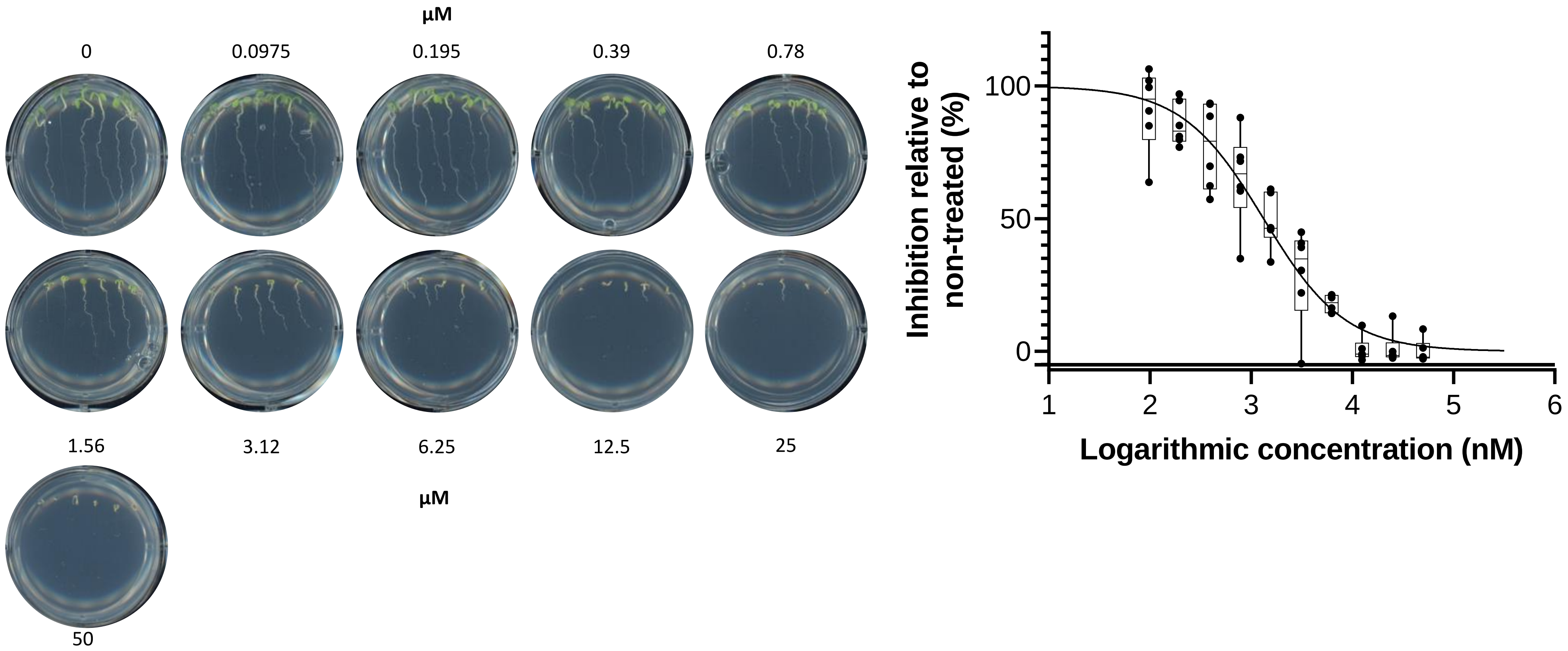

µM
0
0.0975
0.195
0.39
0.78
1.56
3.12
6.25
12.5
25
µM
50
Inhibition relative to non-treated (%)
100
50
0
1
2
3
4
5
6
Logarithmic concentration (nM)

**PJ16**

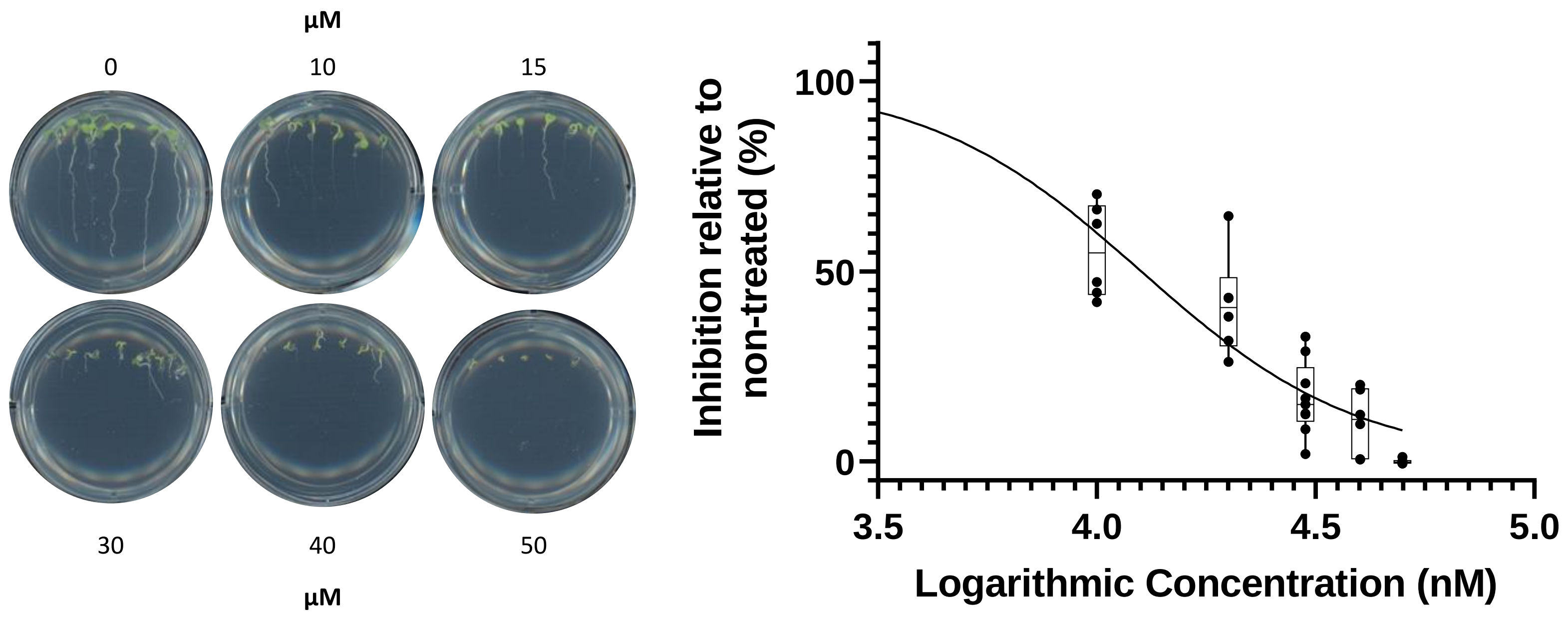
μM
0
10
15
30
40
50
μM
Inhibition relative to non-treated (%)
100
50
0
3.5
4.0
4.5
5.0
Logarithmic Concentration (nM)

**PJ17**

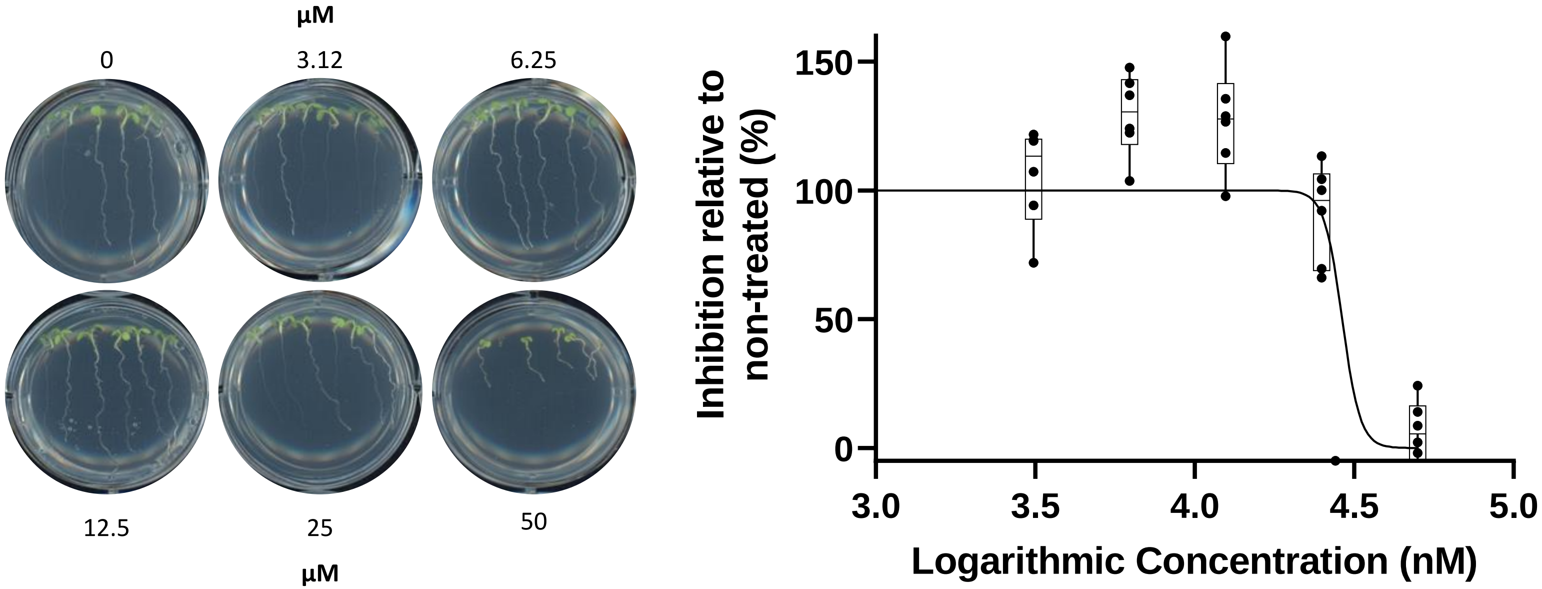
µM
0
3.12
6.25
12.5
25
50
µM
150
100
50
0
3.0
3.5
4.0
4.5
5.0
Inhibition relative to non-treated (%)
Logarithmic Concentration (nM)

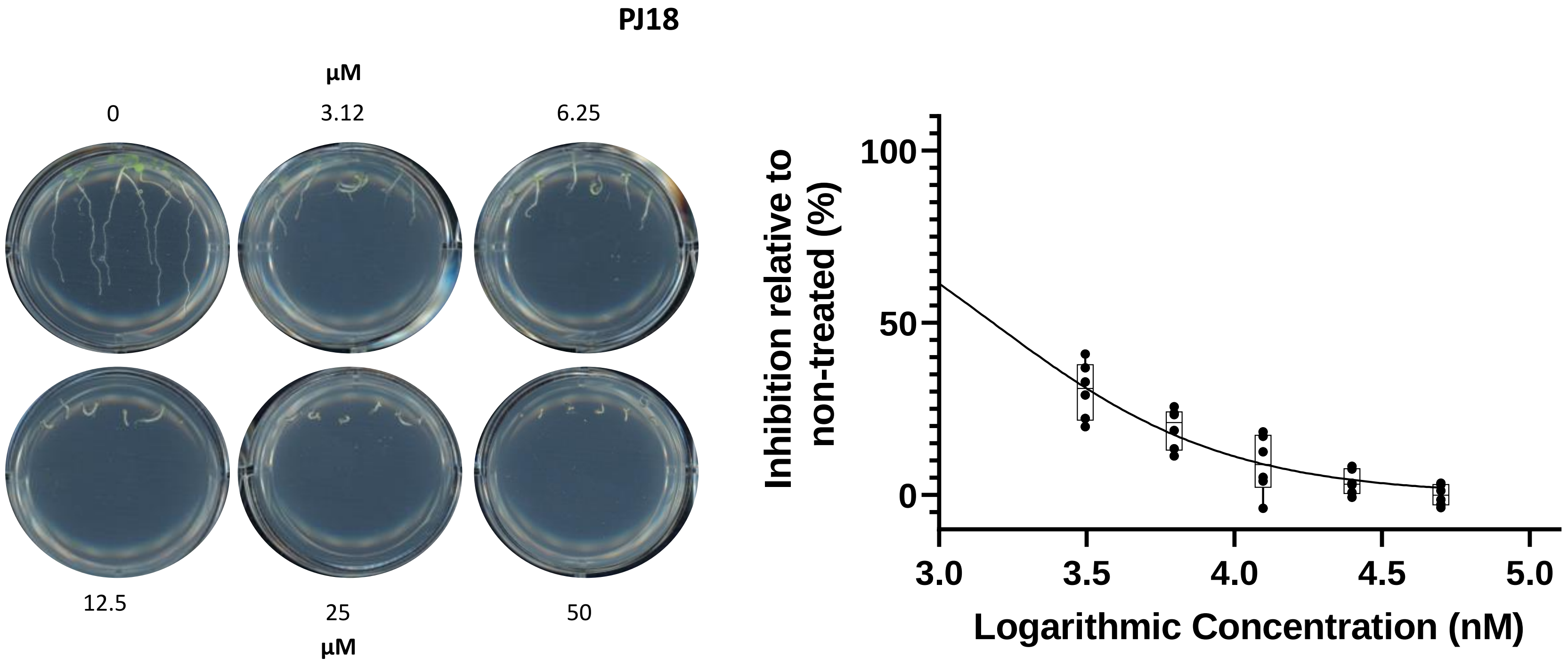
PJ18
µM
0
3.12
6.25
12.5
25
50
µM
100
50
0
Inhibition relative to non-treated (%)
3.0
3.5
4.0
4.5
5.0
Logarithmic Concentration (nM)

**PJ19**

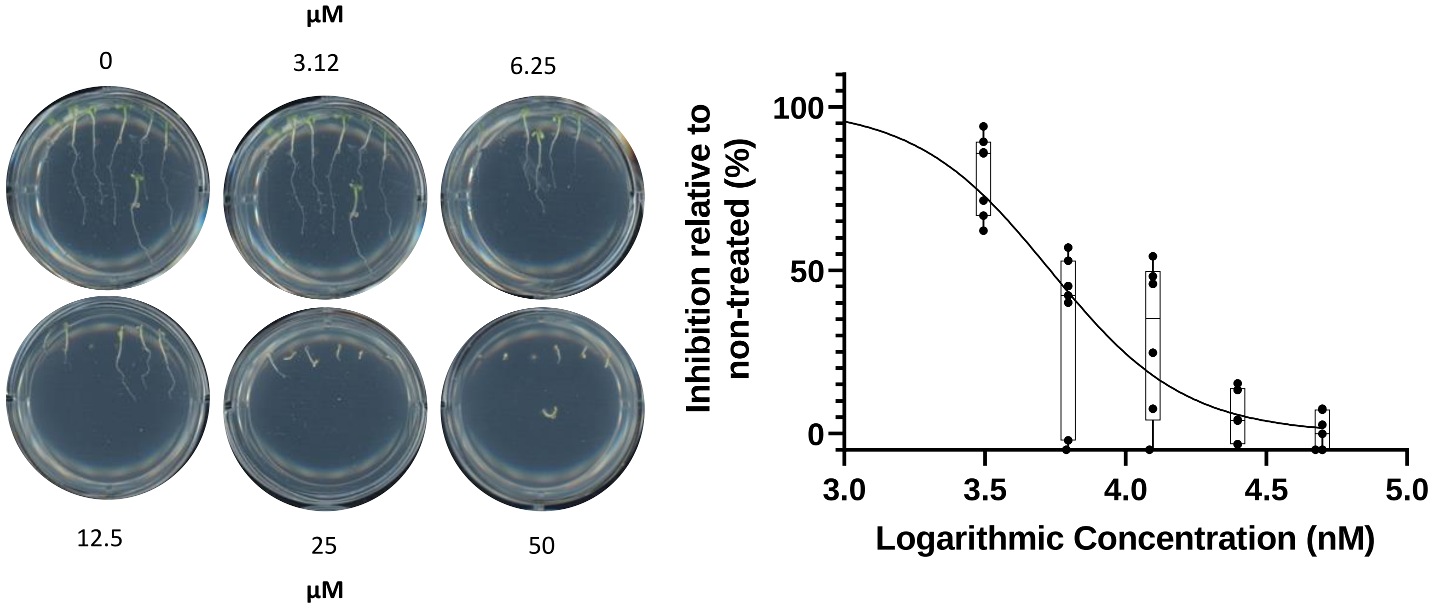

µM
0
3.12
6.25
12.5
25
50
µM
Inhibition relative to non-treated (%)
100
50
0
3.0
3.5
4.0
4.5
5.0
Logarithmic Concentration (nM)

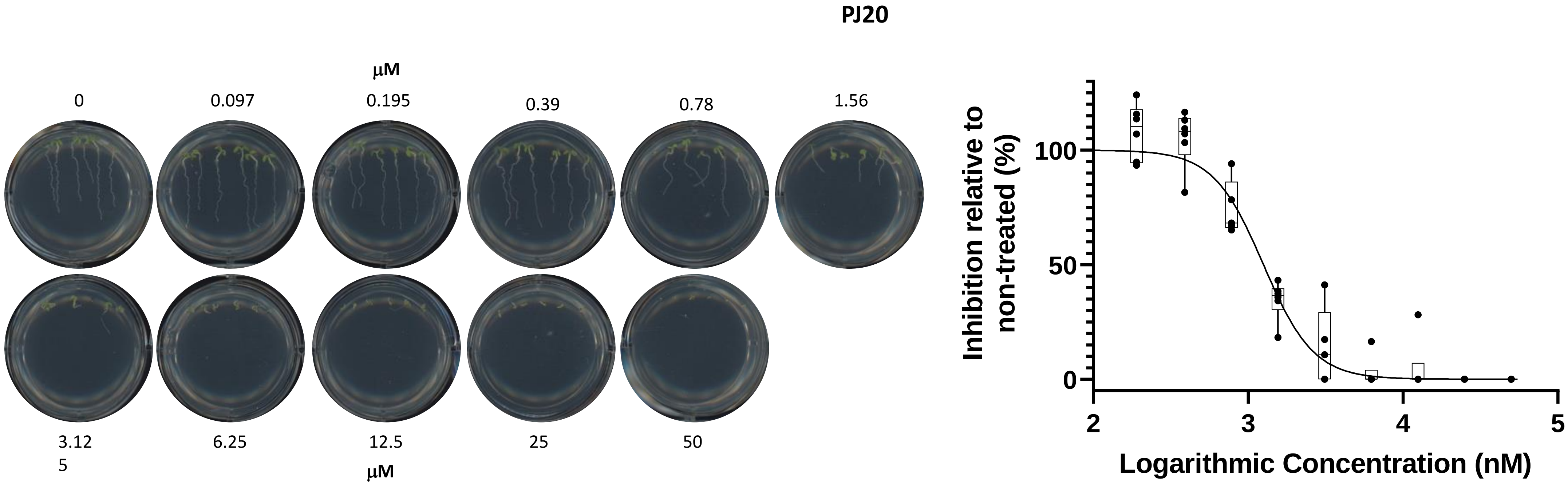
PJ20
μM
0
0.097
0.195
0.39
0.78
1.56
3.12
5
6.25
12.5
25
50
μM
Inhibition relative to non-treated (%)
100
50
0
2
3
4
5
Logarithmic Concentration (nM)

**PJ21**

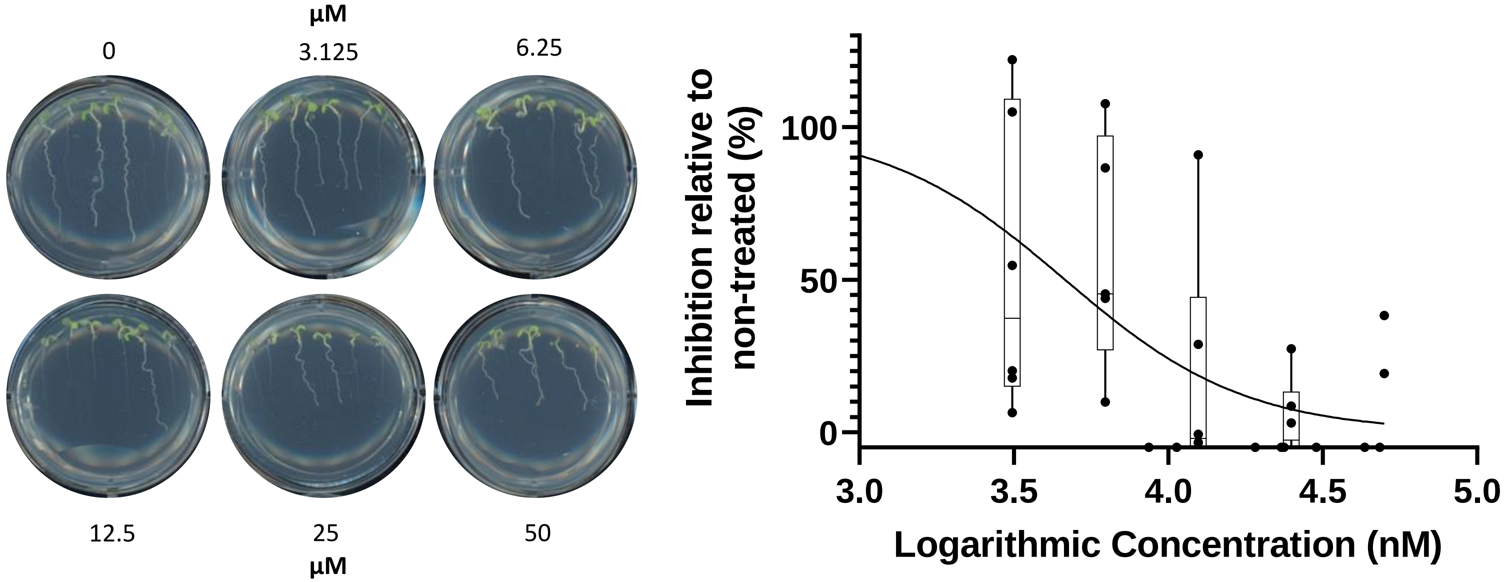

µM
0
3.125
6.25
12.5
25
50
µM
Inhibition relative to non-treated (%)
100
50
0
3.0
3.5
4.0
4.5
5.0
Logarithmic Concentration (nM)

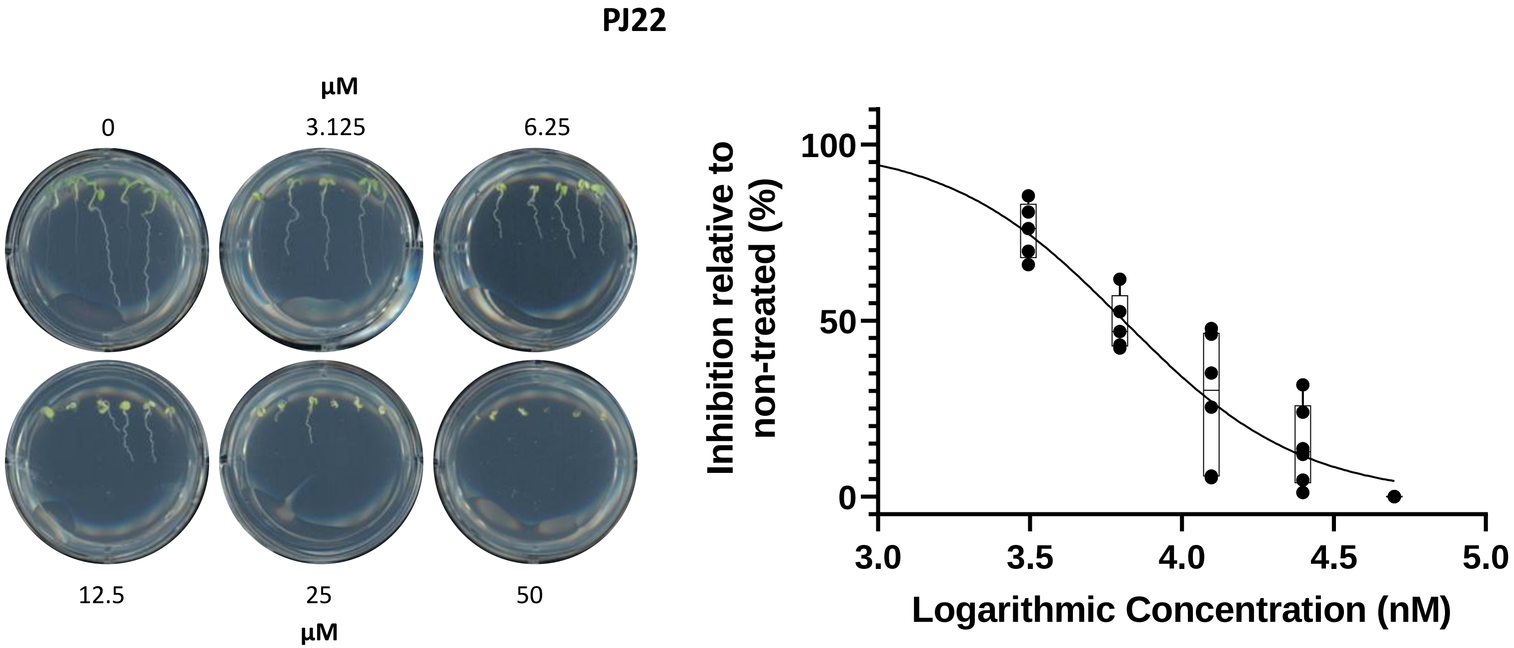

PJ22
µM
0
3.125
6.25
12.5
25
50
µM
Inhibition relative to non-treated (%)
100
50
0
3.0
3.5
4.0
4.5
5.0
Logarithmic Concentration (nM)

**PJ23**

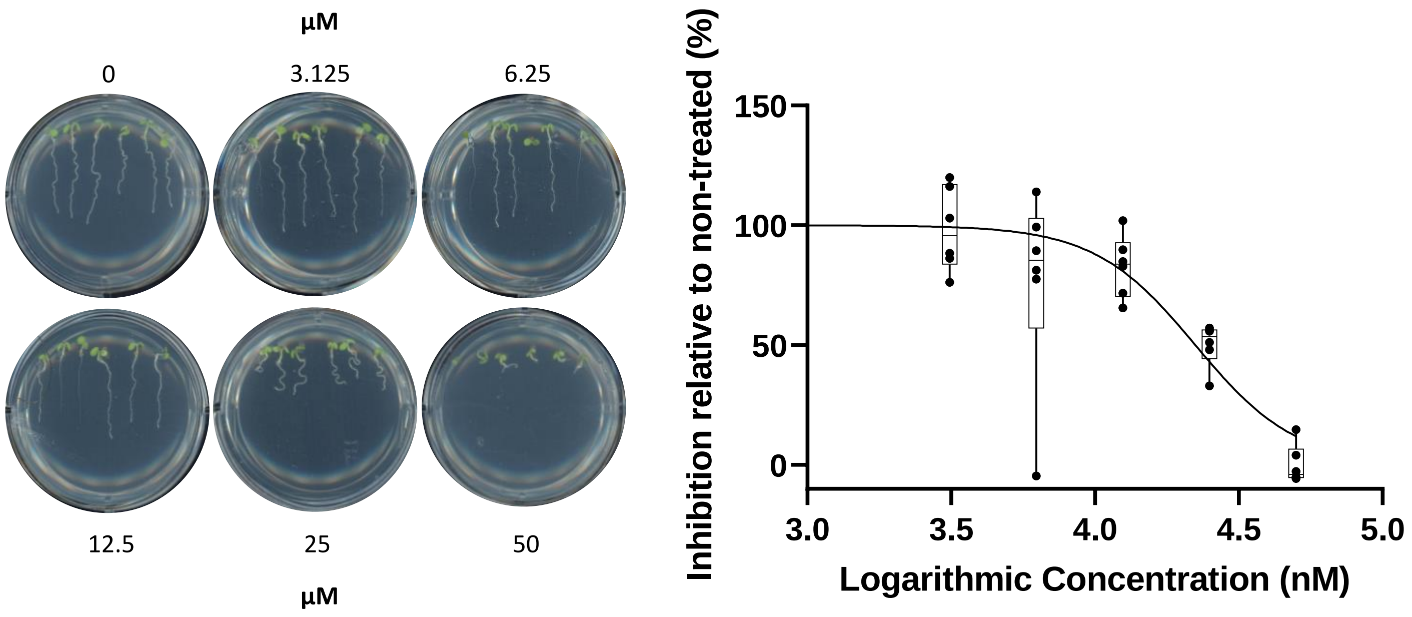
µM
0
3.125
6.25
12.5
25
50
µM
Inhibition relative to non-treated (%)
150
100
50
0
3.0
3.5
4.0
4.5
5.0
Logarithmic Concentration (nM)

# PJ24

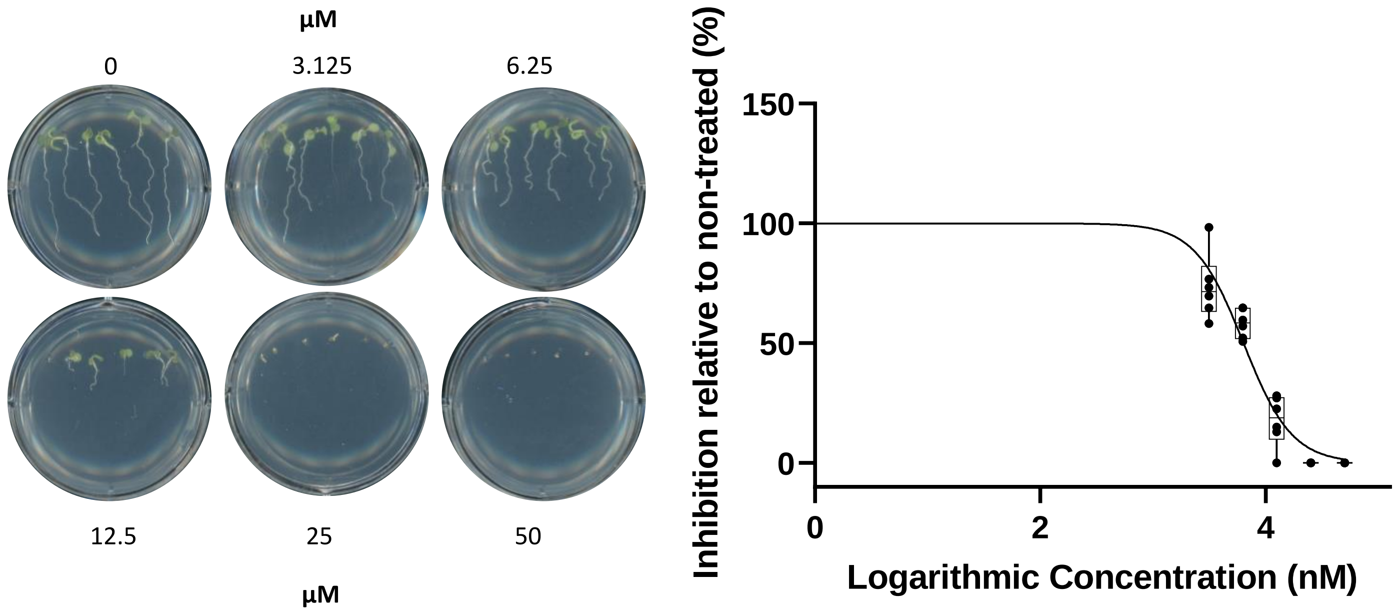
µM
0
3.125
6.25
12.5
25
50
µM
Inhibition relative to non-treated (%)
150
100
50
0
0
2
4
Logarithmic Concentration (nM)

PJ25

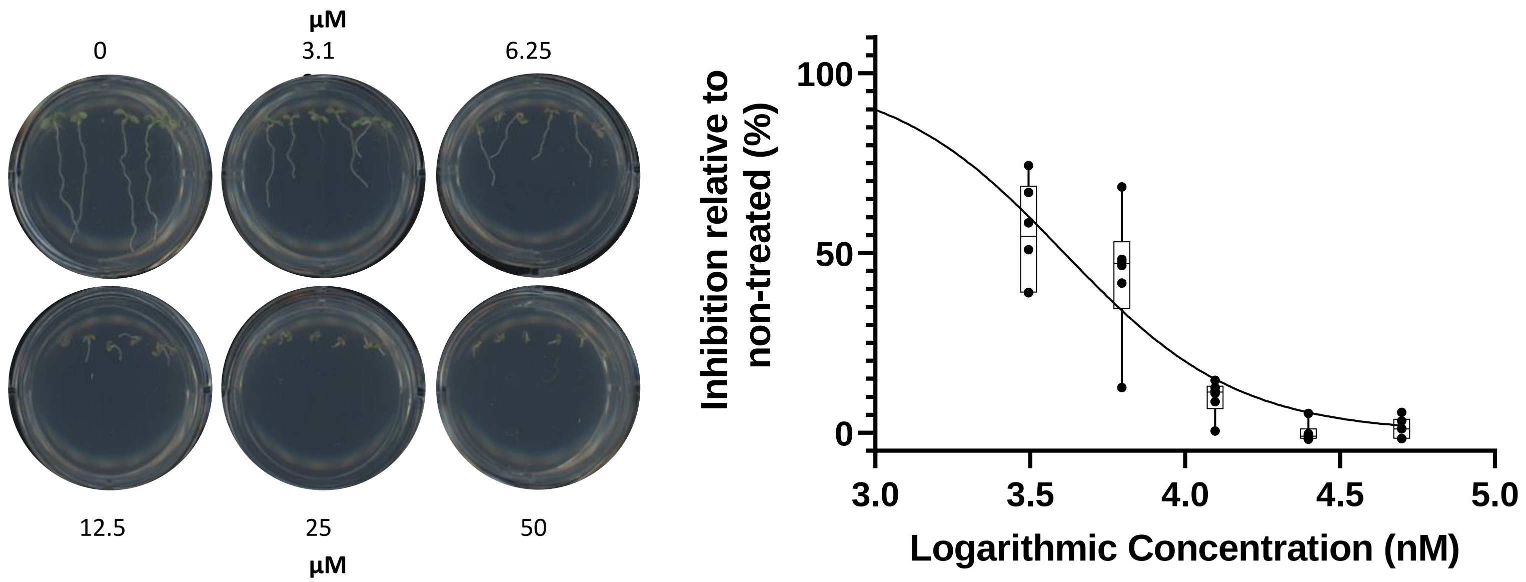

µM
0
3.1
6.25
12.5
25
50
µM
100
50
0
3.0
3.5
4.0
4.5
5.0
Inhibition relative to non-treated (%)
Logarithmic Concentration (nM)

**PJ26**

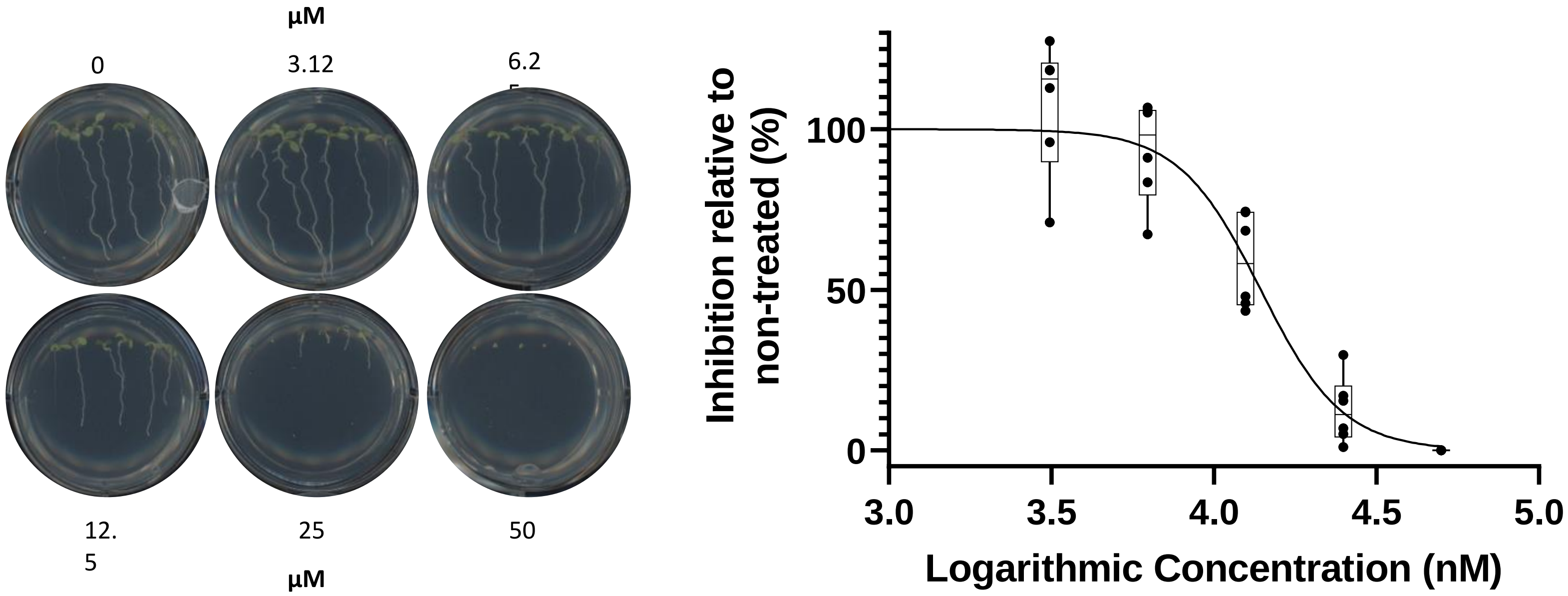
µM
0
3.12
6.2
12.
5
25
50
µM
Inhibition relative to non-treated (%)
100
50
0
3.0
3.5
4.0
4.5
5.0
Logarithmic Concentration (nM)

**PJ29**

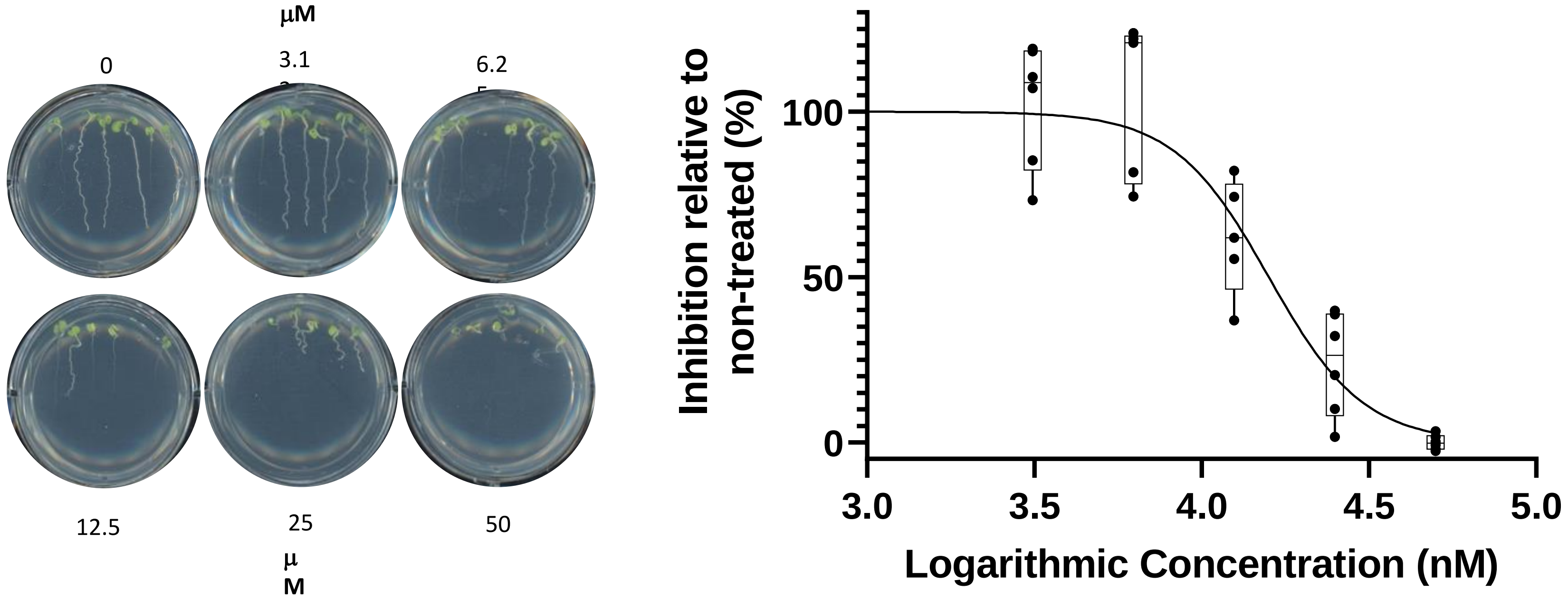
μM
0
3.1
6.2
12.5
25
50
μ
M
Inhibition relative to non-treated (%)
100
50
0
3.0
3.5
4.0
4.5
5.0
Logarithmic Concentration (nM)

**PJ30**

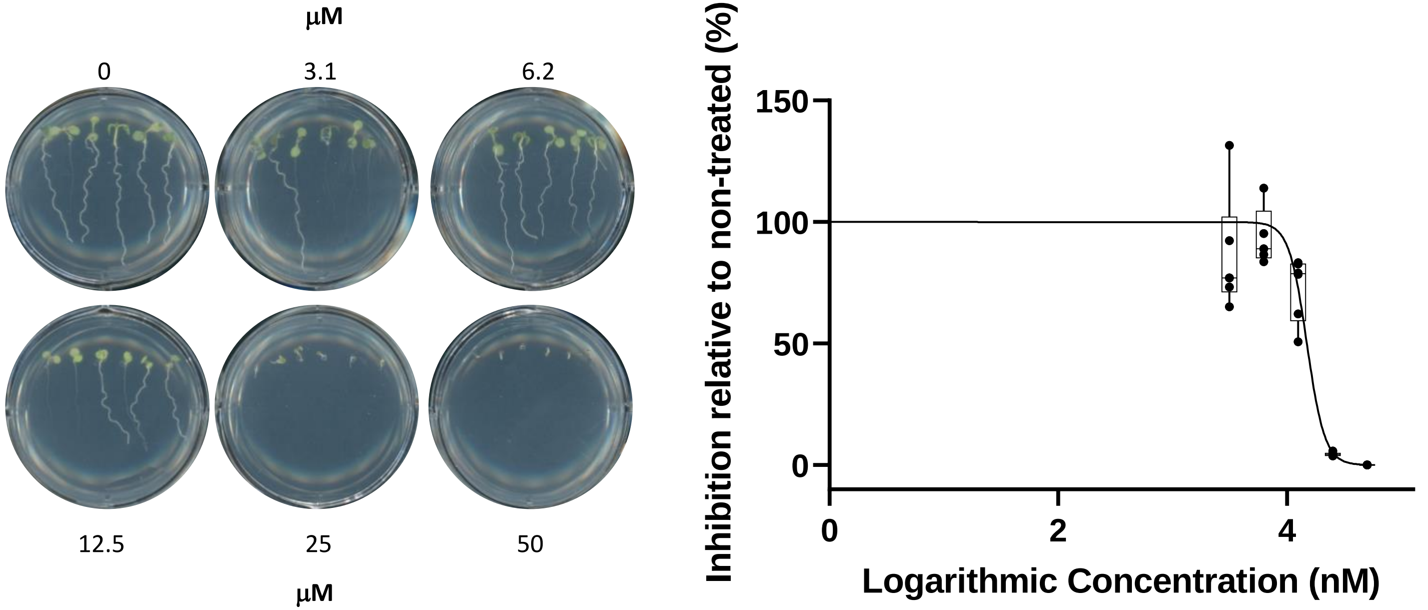

μM
0
3.1
6.2
12.5
25
50
μM
Inhibition relative to non-treated (%)
150
100
50
0
0
2
4
Logarithmic Concentration (nM)

**PJ31**

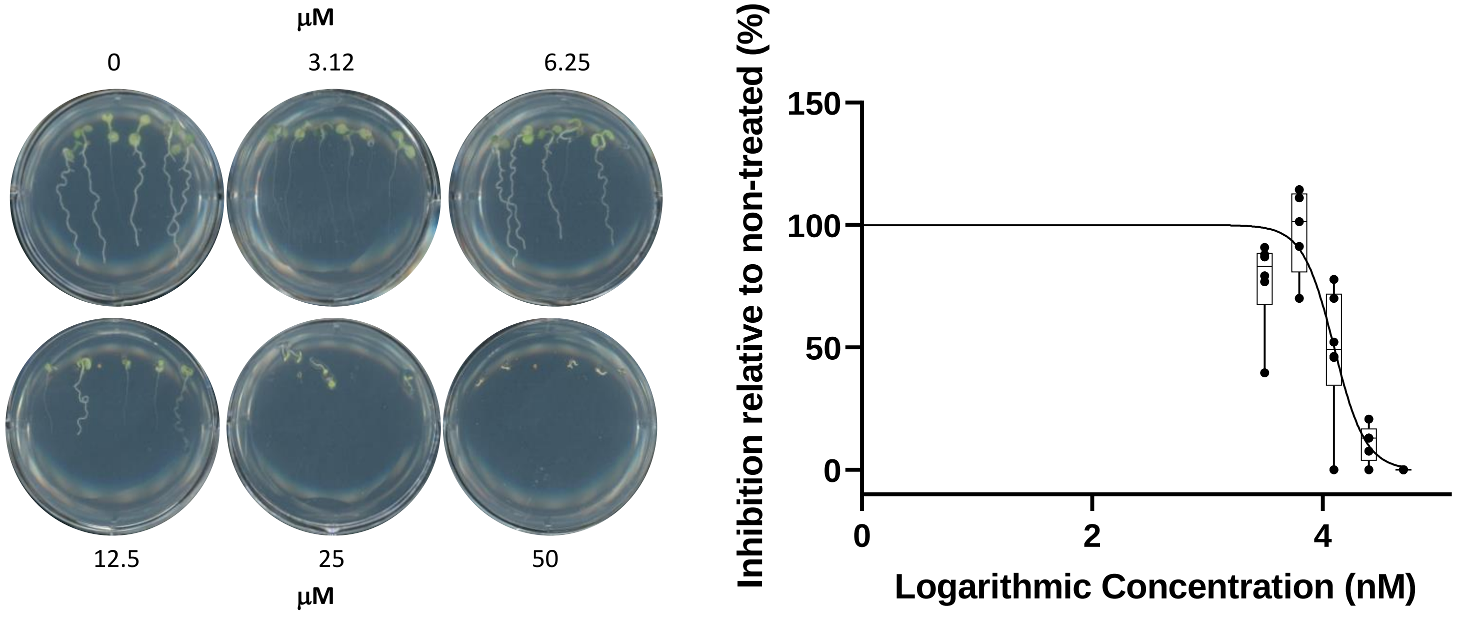

μM
0
3.12
6.25
12.5
25
50
μM
Inhibition relative to non-treated (%)
150
100
50
0
0
2
4
Logarithmic Concentration (nM)

**PJ33**

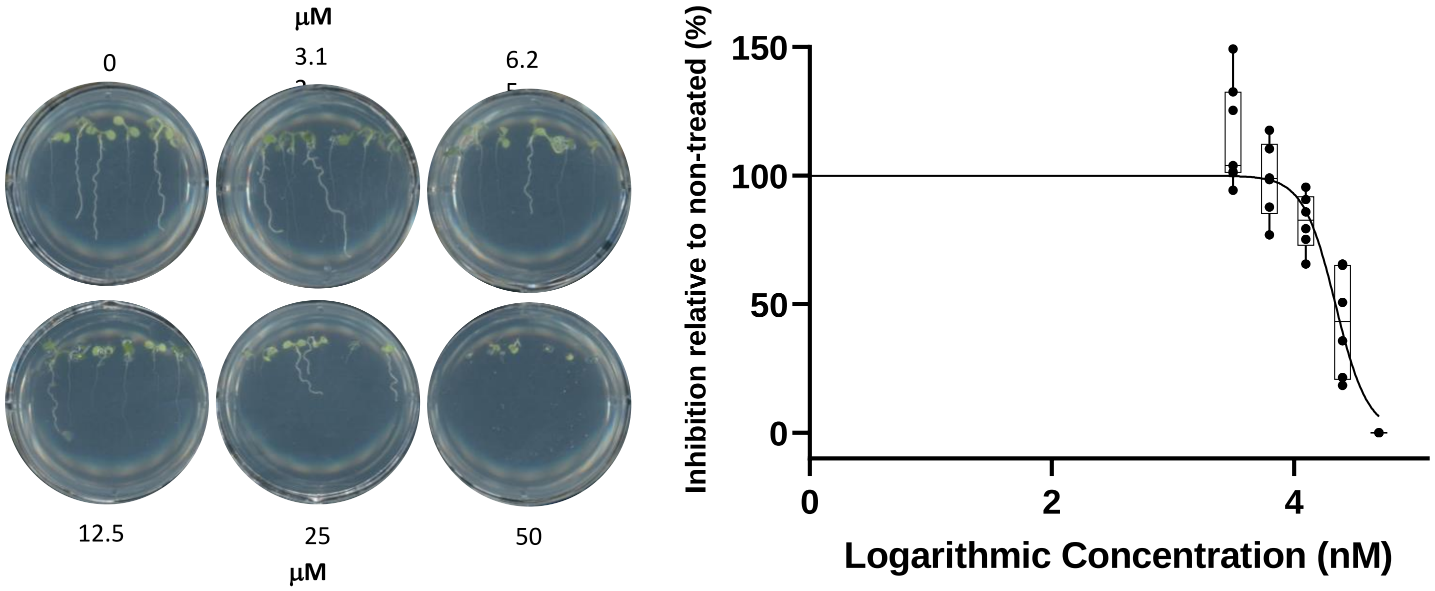
µM
0
3.1
6.2
12.5
25
50
µM
Inhibition relative to non-treated (%)
150
100
50
0
0
2
4
Logarithmic Concentration (nM)

**Basta (Glufosinate)**

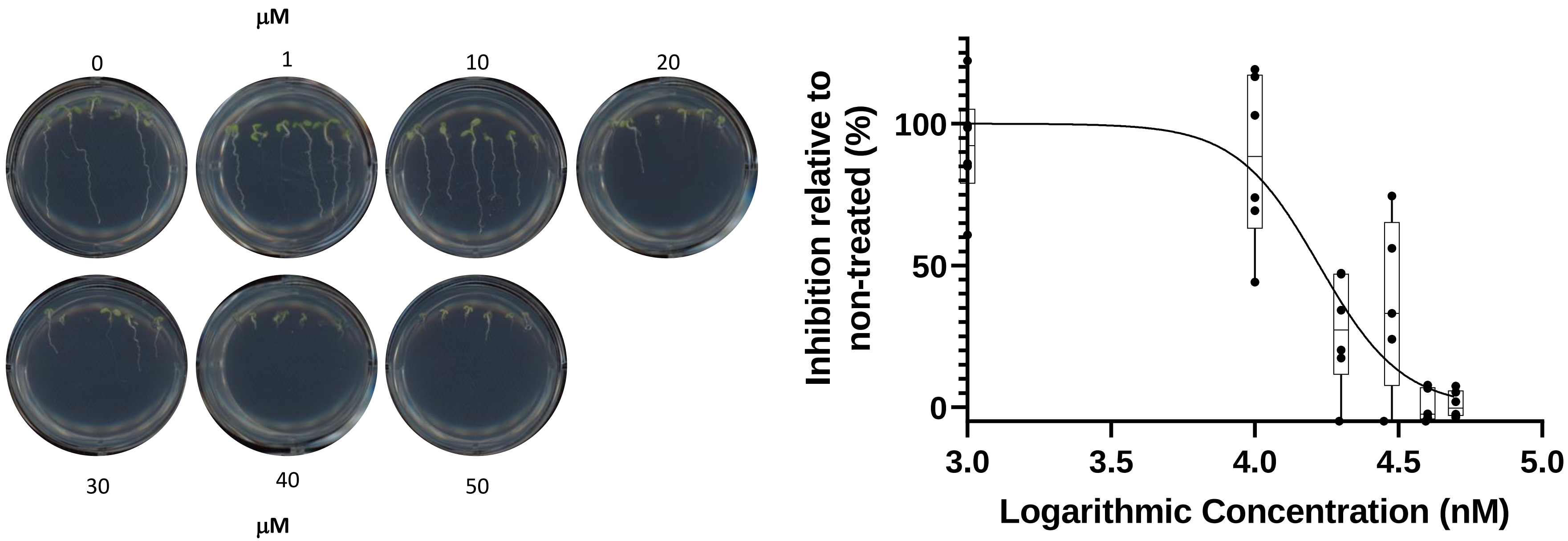
