## Supplementary material for "Pesticide chemical leads inhibiting protein-protein interactions": Table S1

Extended Data, Table 1. Summary of molecules and experimental results

| Molecule assigned name PJ- | M.W (Da) | Canonical_Smiles | FP IC50 (µM) | Arabidopsis roots inhibition@50 µM (% inhibition) | Arabidopsis roots inhibition, IC50 (µM) | Pre emergence clover (% inhibition) | Pre emergence Amaranthus (% inhibition) |
|---|---|---|---|---|---|---|---|
| 1 | 416.469 | CC(C)CNC(=O)c1cccc(NC(=O)c2ccccc2c3ccccc3C(=O)[O-])c1 | 34.4 | --------- | --------- | ND | ND |
| 2 | 415.461 | CC(C)(C)NC(=O)c1cccc(NC(=O)c2ccccc2c3ccccc3C(=O)[O-])c1 | 58.2 | 0 | --------- | ND | ND |
| 3 | 425.413 | [O-]C(=O)c1ccccc1c2ccccc2C(=O)Nc3cccc(NC(=O)c4occc4)c3 | 4.8 | 0 | --------- | ND | ND |
| 4 | 429.468 | COc1ccc(cc1)c2csc(NC(=O)c3ccccc3c4ccccc4C(=O)[O-])n2 | 4.7 | 100 | 9.3 | 88 | 33 |
| 5 | 442.467 | [O-]C(=O)c1ccccc1c2ccccc2C(=O)Nc3cccc(c3)C(=O)Nc4nccs4 | 6.7 | 0 | --------- | ND | ND |
| 6 | 451.450 | Oc1cccc(NC(=O)c2cccc(NC(=O)c3ccccc3c4ccccc4C(=O)[O-])c2)c1 | 6.3 | 0 | --------- | ND | ND |
| 7 | 332.329 | Oc1cccc(NC(=O)c2ccccc2c3ccccc3C(=O)[O-])c1 | 12.9 | 0 | --------- | ND | ND |
| 8 | 432.450 | [O-]C(=O)c1ccccc1c2ccccc2C(=O)Nc3cccc(c3)c4nc5ccccc5[nH]4 | 4.1 | 0 | --------- | ND | ND |
| 9 | 352.387 | CCc1nnc(NC(=O)c2ccccc2c3ccccc3C(=O)[O-])s1 | 30.8 | 0 | --------- | ND | ND |
| 10 | 415.461 | CC(C)(C)NC(=O)c1ccc(NC(=O)c2ccccc2c3ccccc3C(=O)[O-])cc1 | 21.0 | 0 | --------- | ND | ND |
| 11 | 442.467 | [O-]C(=O)c1ccccc1c2ccccc2C(=O)Nc3cccc(NC(=O)c4nccs4)c3 | 10.1 | 0 | --------- | ND | ND |
| 12 | 526.598 | CC=1C=C(C=CC1OCCN2CCOCC2)C=3C=C(NC(=O)C=4C=CC=CC4C=5C=CC=CC5C(=O)O)NN3 | 2.0 | 0 | --------- | ND | ND |
| 13 | 443.530 | NCCC=1C=CC(=CC1)C2=CSC(NC(=O)C=3C=CC=CC3C=4C=CC=CC4C(=O)O)=N2 | 2.0 | 29 | --------- | ND | ND |
| 14 | 455.561 | CC(C)CC=1C=CC(=CC1)C2=CSC(NC(=O)C=3C=CC=CC3C=4C=CC=CC4C(=O)O)=N2 | 4.8 | 17 | 74.0 | ND | ND |
| 15 | 468.343 | OC(=O)C=1C=CC=CC1C=2C=CC=CC2C(=O)NC3=NC(=CS3)C=4C=CC(Cl)=CC4Cl | 5.4 | 91 | 1.3 | 28 | 26 |
| 16 | 444.452 | OC(=O)C=1C=CC=CC1C=2C=CC=CC2C(=O)NC3=NC(=CS3)C=4C=CC(=CC4)[N+](=O)[O-] | 2.6 | 100 | 12.6 | 42 | 0 |
| 17 | 432.458 | NC=1C=C(F)C=CC1C2=CSC(NC(=O)C=3C=CC=CC3C=4C=CC=CC4C(=O)O)=N2 | 2.7 | 72 | 29.0 | ND | ND |
| 18 | 469.331 | OC(=O)C=1C=CC=CC1C=2C=CC=NC2C(=O)NC3=NC(=CS3)C=4C=CC(Cl)=CC4Cl | 12.2 | 100 | 1.5 | 42 | 32 |
| 19 | 463.925 | COC=1C=CC(C2=CSC(NC(=O)C=3C=CC=CC3C=4C=CC=CC4C(=O)O)=N2)=C(Cl)C1 | 3.6 | 100 | 5.4 | 75 | 26 |
| 20 | 459.293 | OC(=O)C=1C=CC=CC1C=2OC=NC2C(=O)NC3=NC(=CS3)C=4C=CC(Cl)=CC4Cl | 248.0 | 100 | 1.3 | 0 | ND |
| 21 | 498.370 | COC=1C=CC(C(=O)O)=C(C1)C=2C=CC=CC2C(=O)NC3=NC(=CS3)C=4C=CC(Cl)=CC4Cl | 6.0 | 50 | 4.6 | 8 | 38 |
| 22 | 486.333 | OC(=O)C=1C=C(F)C=CC1C=2C=CC=CC2C(=O)NC3=NC(=CS3)C=4C=CC(Cl)=CC4Cl | 8.6 | 100 | 6.4 | 63 | 8 |
| 23 | 478.315 | OC(=O)C=1C=CC=CC1C=2C=CC=CC2C(=O)NC3=CC(=CNC3=O)C=4C=CC(Cl)=CC4Cl | 115.0 | 89 | 22.3 | ND | ND |
| 24 | 482.600 | COc1ccc(c2nc(NC(c3c(c4c(C([O-])=O)cccc4)cc(C)cc3)=O)sc2)cc1.[K+] | 3.4 | 100 | 6.3 | 26 | 18 |
| 25 | 482.600 | COc1ccc(c2nc(NC(c3c(c4c(C([O-])=O)cccc4)ccc(C)c3)=O)sc2)cc1.[K+] | 2.1 | 97 | 4.1 | 68 | 24 |
| 26 | 469.556 | [K+].COC=1C=CC(=CC1)C2=CSC(NC(=O)C=3N=CC=CC3C=4C=CC=CC4C(=O)[O-])=N2 | 15.3 | 100 | 13.9 | 2 | 18 |
| 27 | 482.595 | COc1ccc(c2csc(NC(c3cccc(C)c3c4ccccc4C([O-])=O)=O)n2)cc1.[K+] | 18.6 | Non soluble | Non soluble | 55 | ND |
| 28 | 564.478 | [K+].CC(=O)NC=1C=CC(C(=O)NC2=NC(=CS2)C=3C=CC(Cl)=CC3Cl)=C(C1)C=4C=CC=CC4C(=O)[O-] | 11.0 | Non soluble | Non soluble | 0 | ND |
| 29 | 523.425 | [K+].OC=1C=CC(C(=O)NC2=NC(=CS2)C=3C=CC(Cl)=CC3Cl)=C(C1)C=4C=CC=CC4C(=O)[O-] | 3.4 | 98 | 15.9 | 0 | 7 |
| 30 | 537.452 | [K+].OCC=1C=CC(C(=O)NC2=NC(=CS2)C=3C=CC(Cl)=CC3Cl)=C(C1)C=4C=CC=CC4C(=O)[O-] | 9.4 | 100 | 14.8 | 37 | 87 |
| 31 | 523.425 | [K+].OC=1C=CC(=C(C1)C(=O)NC2=NC(=CS2)C=3C=CC(Cl)=CC3Cl)C=4C=CC=CC4C(=O)[O-] | 4.5 | 95 | 12.5 | 0 | 0 |
| 32 | 522.441 | [K+].NC=1C=CC(C(=O)NC2=NC(=CS2)C=3C=CC(Cl)=CC3Cl)=C(C1)C=4C=CC=CC4C(=O)[O-] | 7.7 | 100 | 18.0 | 88 | 88 |
| 33 | 482.600 | COc1ccc(c2c(C)sc(NC(c3c(c4c(C([O-])=O)cccc4)cccc3)=O)n2)cc1.[K+] | 6.7 | 72 | 21.0 | 0 | 0 |
| 34 | 495.580 | COc1ccc(c2nn(CCO)c(NC(c3c(c4c(C([O-])=O)cccc4)cccc3)=O)c2)cc1.[K+] | 49.5 | ------------- | ------------- | 41 | ND |
| 35 | 459.520 | O=C(C1=C(C2=C(C(NC3=NC(C4=CC=C(OC)C=C4)=CS3)=O)N=CO2)C=CC=C1)[O-].[K+] | 4587.0 | 86 | ------------- | 12 | 0 |
